# 50S subunit recognition and modification by the *Mycobacterium tuberculosis* ribosomal RNA methyltransferase TlyA

**DOI:** 10.1101/2021.11.11.467980

**Authors:** Zane T. Laughlin, Suparno Nandi, Debayan Dey, Natalia Zelinskaya, Marta A. Witek, Pooja Srinivas, Ha An Nguyen, Emily G. Kuiper, Lindsay R. Comstock, Christine M. Dunham, Graeme L. Conn

## Abstract

Changes in bacterial ribosomal RNA methylation status can alter the activity of diverse groups of ribosome-targeting antibiotics. These modifications are typically incorporated by a single methyltransferase that acts on one nucleotide target and rRNA methylation directly prevents drug binding, thereby conferring drug resistance. Loss of intrinsic methylation can also result in antibiotic resistance. For example, *Mycobacterium tuberculosis* becomes sensitized to tuberactinomycin antibiotics, such as capreomycin and viomycin, due to the action of the intrinsic methyltransferase TlyA. TlyA is unique among antibiotic resistance-associated methyltransferases as it has dual 16S and 23S rRNA substrate specificity and can incorporate cytidine-2’-O-methylations within two structurally distinct contexts. Here, we report the structure of a mycobacterial 50S subunit-TlyA complex trapped in a post-catalytic state with a S-adenosyl-L-methionine analog using single-particle cryogenic electron microscopy. Together with complementary functional analyses, this structure reveals critical roles in 23S rRNA substrate recognition for conserved residues across an interaction surface that spans both TlyA domains. These interactions position the TlyA active site over the target nucleotide C2144 which is flipped from 23S Helix 69 in a process stabilized by stacking of TlyA residue Phe157 on the adjacent A2143. Base flipping may thus be a common strategy among rRNA methyltransferase enzymes even in cases where the target site is accessible without such structural reorganization. Finally, functional studies with 30S subunit suggest that the same TlyA interaction surface is employed to recognize this second substrate, but with distinct dependencies on essential conserved residues.

**Significance Statement:** The bacterial ribosome is an important target for antibiotics used to treat infection. However, resistance to these essential drugs can arise through changes in ribosomal RNA (rRNA) modification patterns through the action of intrinsic or acquired rRNA methyltransferase enzymes. How these antibiotic resistance-associated enzymes recognize their ribosomal targets for site-specific modification is currently not well defined. Here, we uncover the molecular basis for large ribosomal (50S) subunit substrate recognition and modification by the *Mycobacterium tuberculosis* methyltransferase TlyA, necessary for optimal activity of the antitubercular drug capreomycin. From this work, recognition of complex rRNA structures distant from the site of modification and “flipping” of the target nucleotide base both emerge as general themes in ribosome recognition for bacterial rRNA modifying enzymes.

## Introduction

Ribosome-targeting antibiotics are a structurally and mechanistically diverse group of anti-infectives that comprise a significant proportion of currently used treatments for bacterial infections (1, 2). However, among resistance mechanisms exploited by pathogenic bacteria to evade the effects of these antibiotics, ribosomal RNA (rRNA) drug-binding site methylation is already established or is quickly emerging as a major threat to such treatments (3, 4). For example, diverse human pathogens have acquired resistance modifications that impact the efficacy of aminoglycosides, macrolides and multiple other drug classes targeting the ribosome peptidyl transferase center. These modifications are incorporated by S-adenosyl-L-methionine (SAM)-dependent methyltransferases such as the Class I aminoglycoside-resistance 16S rRNA methyltransferases (e.g. NpmA and ArmA/RmtA-H), the Class I Erm family methyltransferases, and the radical SAM enzyme Cfr (5–7). Less commonly, reduced intrinsic methylation can also lead to resistance, such as for kasugamycin, streptomycin or capreomycin through loss of activity of the 16S rRNA methyltransferases RsmG/GidB (8, 9), KsgA (10), and TlyA (11), respectively.

Capreomycin is a member of the tuberactinomycin class of ribosome-targeting antibiotics and has an important history in the treatment of *Mycobacterium tuberculosis* (*Mtb*) infections resistant to the first-line drugs rifampin and isoniazid (12). Capreomycin binds at the subunit interface of mature 70S ribosomes, adjacent to 16S rRNA helix 44 (h44) of the small (30S) subunit and 23S rRNA Helix 69 (H69) of the large (50S) subunit (13). In a recent study, the tuberactinomycin antibiotic viomycin was also found to bind the 70S ribosome at several other locations (14), suggesting that this class of antibiotics may target multiple ribosomal sites to interfere with translation. Capreomycin’s activity is thought to arise via stabilization of tRNA in the A site of the ribosome, thereby halting translation (13). Capreomycin has also been proposed to disrupt the interaction of ribosomal proteins uL10 and bL12, thereby blocking binding of elongation factors during translation (15). However, this mechanism is harder to reconcile with the binding sites of capreomycin and viomycin which are distant from both uL10 or bL12 (13, 14, 16), as well as the impact of changes in rRNA modification status in the A site on their activity.

Capreomycin binding to the *Mtb* 70S ribosome is dependent on 2’-O methylation of two nucleotides at the subunit interface, 16S rRNA C1392 and 23S rRNA C2144 (**Fig. 1A**; corresponding to *E. coli* nucleotides C1409 and C1920, respectively) (17). While the precise role of these modifications in ribosome structure and function is currently unclear, it is thought that they may somehow change the conformation of the rRNA allowing for optimal capreomycin binding (17, 18). Evolutionary maintenance of intrinsic rRNA modifications which increase sensitivity to antibiotics may be driven by their contribution to optimal fitness in the absence of drug or through decreased stability of unmodified 70S, as observed in *Mycolicibacterium smegmatis* (formerly *Mycobacterium smegmatis*; *Msm*) and *Campylobacter jejuni*, respectively (19, 20). Both modifications are incorporated by a single SAM-dependent ribose 2’-O-methyltransferase, TlyA, encoded by Rv1694 in *Mtb* (18). TlyA has strong substrate preference for intact ribosomal subunits over free 16S or 23S rRNA, and individual modification of isolated subunits occurs prior to 70S assembly due to the target site locations on the interface surfaces of their respective subunits (**Fig. 1A**) (17). The TlyA family of methyltransferases is also divided into two groups based on their substrate specificities: Type I TlyA (TlyA^I^) exclusively methylate 23S rRNA, while the slightly larger TlyA^II^, including the *Mtb* enzyme, possess dual 16S and 23S specificity (17). However, how *Mtb* TlyA and other TlyA^II^ enzymes recognize and modify these two structurally distinct substrates is not currently known.

**Figure 1.**
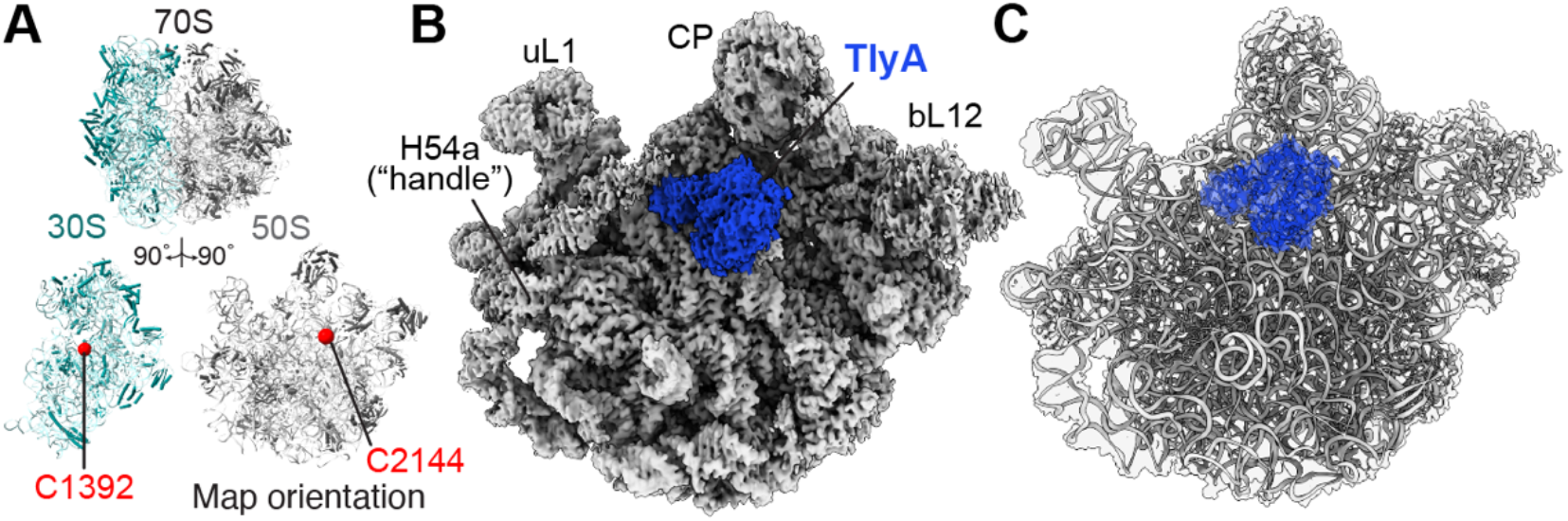
Cryo-EM map at 3.05 Å resolution of the 50S-TlyA complex. ***A,*** Cartoon of the *Msm* 70S ribosome (*top*) and individual 30S and 50S subunits rotated to show their intersubunit interface surfaces (*bottom*), with the nucleotides modified by TlyA indicated (red spheres). ***B,*** Final overall cryo-EM map for the 50S-TlyA complex. TlyA (blue) is bound to the 50S subunit (white) on the subunit interface over H69 containing the modification site (residue C2144). Key 50S subunit features are indicated: uL1 stalk (uL1), central protuberance (CP), the bL12 stalk (bL12) and H54a (also known as the handle). H54a, which extends outward and interacts extensively with 30S subunit in the intact 70S ribosome, was partially observed on the 50S subunit surface in proximity to TlyA, but with weaker map features close to the enzyme. ***C*,** Final model of the 50S-TlyA complex shown with semi-transparent map (white behind the model for 50S and blue in front of the model for TlyA).

We previously determined the crystal structure of the C-terminal domain (CTD) of *Mtb* TlyA with and without a four amino acid interdomain linker sequence (21). The TlyA CTD adopts the expected Class I methyltransferase fold but was unexpectedly found to be deficient in SAM binding in the absence of the interdomain linker. A TlyA CTD structure including the linker also revealed that this short motif can either extend the first α-helix of the CTD, or form a loop structure similar to that proposed earlier via homology modeling (21, 22). While a structure of the TlyA N-terminal domain is currently not available, modeling suggests a ribosomal protein S4-like domain (22). Collectively, these findings suggest that the N-terminal domain may be essential for rRNA recognition and binding, with the interdomain linker potentially playing a role in promoting SAM binding and methyltransferase activity in the CTD once bound to the correct substrate (21).

Here, we describe the structure of full-length *Mtb* TlyA bound to the *Msm* 50S subunit (hereafter, 50S-TlyA). The structure reveals the critical role played by the TlyA NTD in recognizing a complex 23S rRNA structure at the base of H69, positioning the TlyA CTD on H69 with its active site over the target nucleotide, C2144 (in *Msm* numbering, which is used exclusively hereafter unless noted). In addition, we find that TlyA uses a mechanism of base flipping for target site recognition and modification despite the accessibility of the C2144 ribose 2’-OH in H69, suggesting that this may be a general strategy for substrate molecular recognition among rRNA methyltransferases.

## Results

### Determination of the 50S-TlyA complex structure

50S subunits without ribose modification on 23S rRNA nucleotide C2144 were isolated from *Msm* strain LR222 C101A which lacks TlyA activity and *Mtb* TlyA was expressed in *E. coli* and purified as previously described (21). A SAM-analog, “N-mustard 6” (NM6), was used to increase occupancy of TlyA on the 50S subunit (23, 24); NM6 is transferred by TlyA in its entirety to the ribose 2’-OH of C2144 and its covalent attachment to the 23S rRNA thus stabilizes the 50S-TlyA complex by virtue of the enzyme’s affinity for both substrate and SAM analog (**Fig. S1**). Using this approach, we determined a 3.05 Å-resolution overall map of TlyA bound to the 50S subunit in a state immediately after catalysis of C2144 ribose modification by single-particle cryogenic electron microscopy (cryo-EM) (**Fig. 1; Fig. S2, Fig. S3A-C**).

One 23S rRNA feature, H54a (also called the “handle”), was significantly shifted from its position in the previously solved *Msm* 70S structure where it makes extensive interactions with the 30S subunit (PDB code 5O60) (25). This feature was visible in some 3D reconstructions (**Fig. S4**), where H54a lies across the subunit interface surface of the 50S subunit. In contrast, in other reconstructions, the map was weaker, suggesting H54a is dynamic in the free 50S subunit. H54a’s variable position and weak map adjacent to the bound TlyA in most 3D reconstructions suggest that H54a does not contribute to TlyA interaction with the 50S subunit despite its proximity to the enzyme NTD (**Fig. S4**).

An initial model for the 50S-TlyA complex was produced by docking a model of full-length TlyA, previously produced using a combination of NTD homology model and CTD crystal structure (PDB code 5KYG) (21), into unoccupied map surrounding 23S rRNA H69. This TlyA structure was subsequently rebuilt in Coot (26), including a complete rebuilding of the NTD (see Methods and Materials). Although the map was of sufficient quality for initial rebuilding of the NTD, the region corresponding to the TlyA CTD was less well defined. We therefore also performed multibody refinement with H69 and TlyA masked to separate this specific region of interest from the remainder of the 50S subunit (**Fig. S2**). This multibody refinement produced separate maps of the H69:TlyA complex (3.61 Å following post-processing; **Figs. S3D-F**) and the remaining 50S subunit structure lacking H69 (2.99 Å following post-processing; **Figs. S3G-I**). The former map was significantly improved compared to the corresponding region of the original map, with more information on the secondary structure and side chains of TlyA providing insights into how full-length TlyA interacts with its 23S rRNA substrate. Each map from multibody refinement was used for final model building and refinement of its associated structure, and the separate structures combined to generate a complete model of the 50S-TlyA complex (**Fig. 2; Fig. S2**).

**Figure 2.**
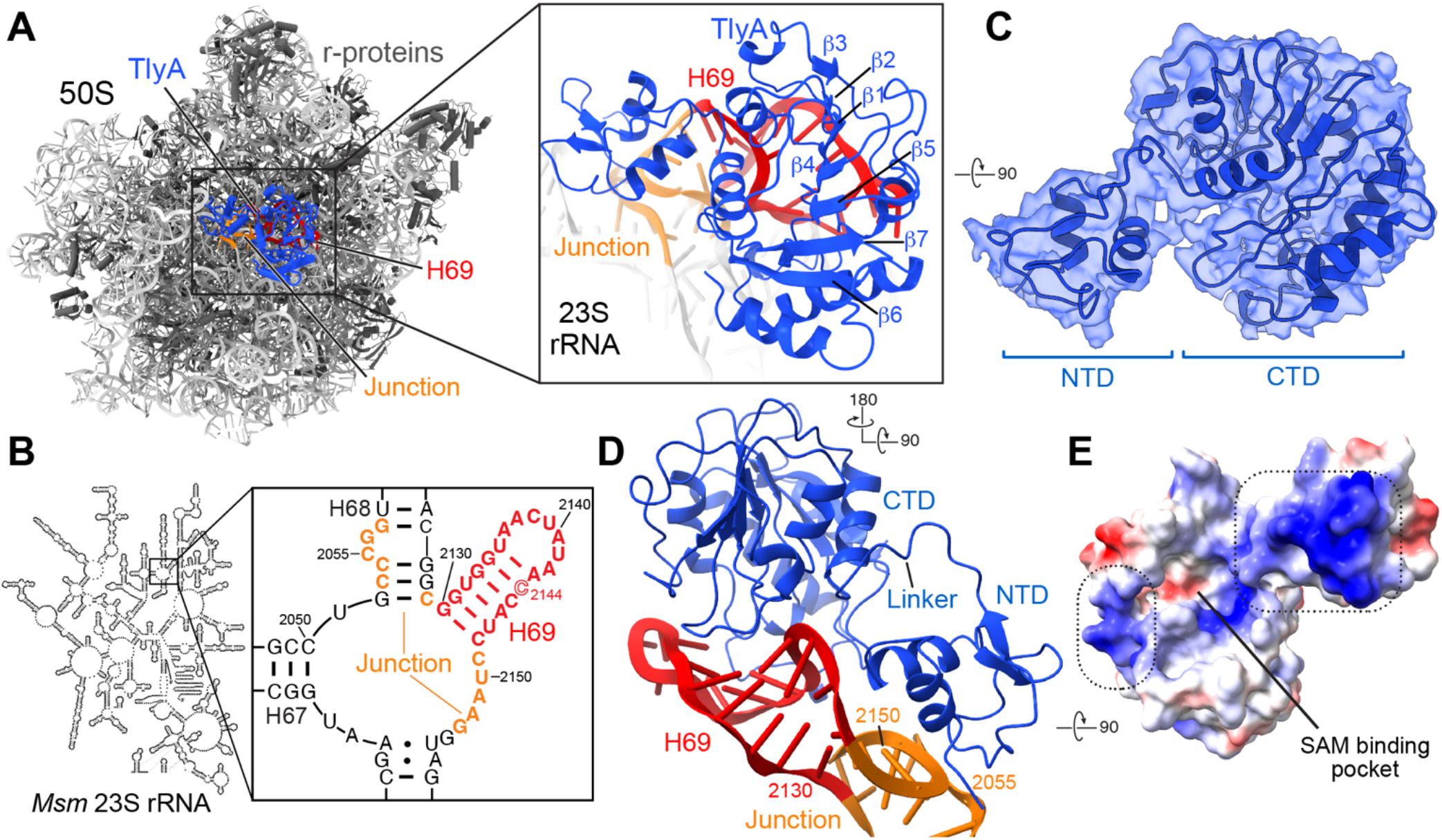
TlyA binds to 23S rRNA H69 and the adjacent rRNA junction via a surface of positively charged residues. ***A***, Structure of the 50S-TlyA complex with TlyA (blue cartoon) bound at H69 (red) and adjacent junction (orange) on the 50S subunit interface surface. Ribosomal proteins are shown in dark gray and the remaining 23S rRNA in white. *Inset*, zoomed-in view of TlyA bound to H69 and the adjacent rRNA junction. ***B***, *Msm* 23S rRNA secondary structure, highlighting the sequence of regions bound by TlyA: H69 and the adjacent junction. ***C***, Modeled structure of full-length TlyA comprising an N-terminal ribosomal protein S4 fold (NTD) and a C-terminal Class I methyltransferase fold with a seven β-strand core (β1-β7, labeled in *panel A*) surrounded by α-helices. The structure is shown in an orthogonal view to *panel A* with surrounding multibody map. ***D***, The TlyA-H69/junction interaction viewed from the 50S subunit surface. The TlyA NTD binds at the base of H69 and the adjacent 23S rRNA junction, while the TlyA CTD interacts exclusively with H69 nucleotides around the modification site. ***E***, Electrostatic surface of TlyA revealing two main patches of positively charged surface (blue, dashed boxes) along the face of TlyA interacting with the rRNA. The negatively charged (red) cosubstrate-binding pocket is indicated between the two positive patches.

When bound to the 50S subunit, the TlyA CTD is essentially identical to the previously determined structure of the isolated domain (PDB code 5KYG; 2.45 Å RMSD for 209 CTD Cα atoms), with the exception of a significant movement (~6-8 Å) of the loop containing residues 114-117 that is necessary to avoid clash with the minor groove surface of H69 (**Fig. S5A,B**). TlyA is structurally similar to the putative *S. thermophilus* hemolysin (PDB code 3HP7), but with a significant difference in the relative NTD and CTD orientation as a result of the distinct backbone path at the linker between the domains (**Fig. S5C,D**). The structures thus align well for superpositions based on either individual domain (3.68 and 2.00 Å RMSD for 209 CTD or 59 NTD Cα atoms, respectively), but less well for the full protein (overall 7.13 Å RMSD for 268 Cα atoms). The final NTD model reveals a globular domain with expected similarity to ribosomal protein S4 (2.55 Å RMSD for 59 NTD Cα atoms; **Fig. S5E**), comprising two adjacent short α-helices (residues 6-14 and 20-28) and two short β-strands (residues 32-24 and 52-54) preceding an interdomain linker (residues 60-63).

As described further in the following sections, TlyA binds the 50S subunit on its subunit interaction surface, with both TlyA domains surrounding H69 and the NTD making additional contacts to the rRNA junction at the base of H69 (**Fig. 2A-D**). Together, the two TlyA domains form a continuous positively charged surface in contact with the 23 rRNA, suggesting that both play an important role in 50S subunit binding and specific substrate recognition (**Fig. 2D,E**). The final model also reveals the CTD of TlyA with bound SAM analog NM6 positioned directly over H69 residue C2144 in a post-catalytic state (i.e., with C2144 modified with NM6).

### TlyA NTD residues Arg6 and Arg20 exploit a complex rRNA structure for specific substrate recognition

Eight residues in the TlyA NTD were identified to make potentially critical interactions with nucleotides at the base of H69 and the adjacent rRNA junction: Arg4, Arg6, Arg18, Ser19, Arg20, Gln21, Gln22, and Lys41 (**Fig. 3A**). Three of these residues, Arg4, Arg6 and Arg20, are clustered around a complex (non-A-form helical) RNA structure formed by nucelotides C2149-G2153 of the 23 rRNA sequence immediately following H69 (**Fig. 3B; Fig. S6A**). While Arg4 is positioned to form a single electrostatic interaction with the phosphate group of A2151, Arg6 and Arg20 each form interaction networks with multiple rRNA nucleotides and with each other, likely stabilized by additional interactions with Asp8 (**Fig. 3B**). Specifically, Arg6 recognizes a sharp turn in the rRNA backbone via contacts with the bridging oxygen of A2151 and non-bridging oxygens of A1552, as well as a cation-π stacking interaction on the nucleobase of U2150. Similarly, Arg20 recognizes the phosphate backbone of 23S rRNA via electrostatic interactions with the phosphate groups of C2149 and U2150 (**Fig. 3B**). Consistent with critical roles in 23S rRNA recognition for Arg6 and Arg20, these two residues, as well as Asp8, are almost universally conserved among TlyA homologs (**Fig. 3C; Fig. S7**).

**Figure 3.**
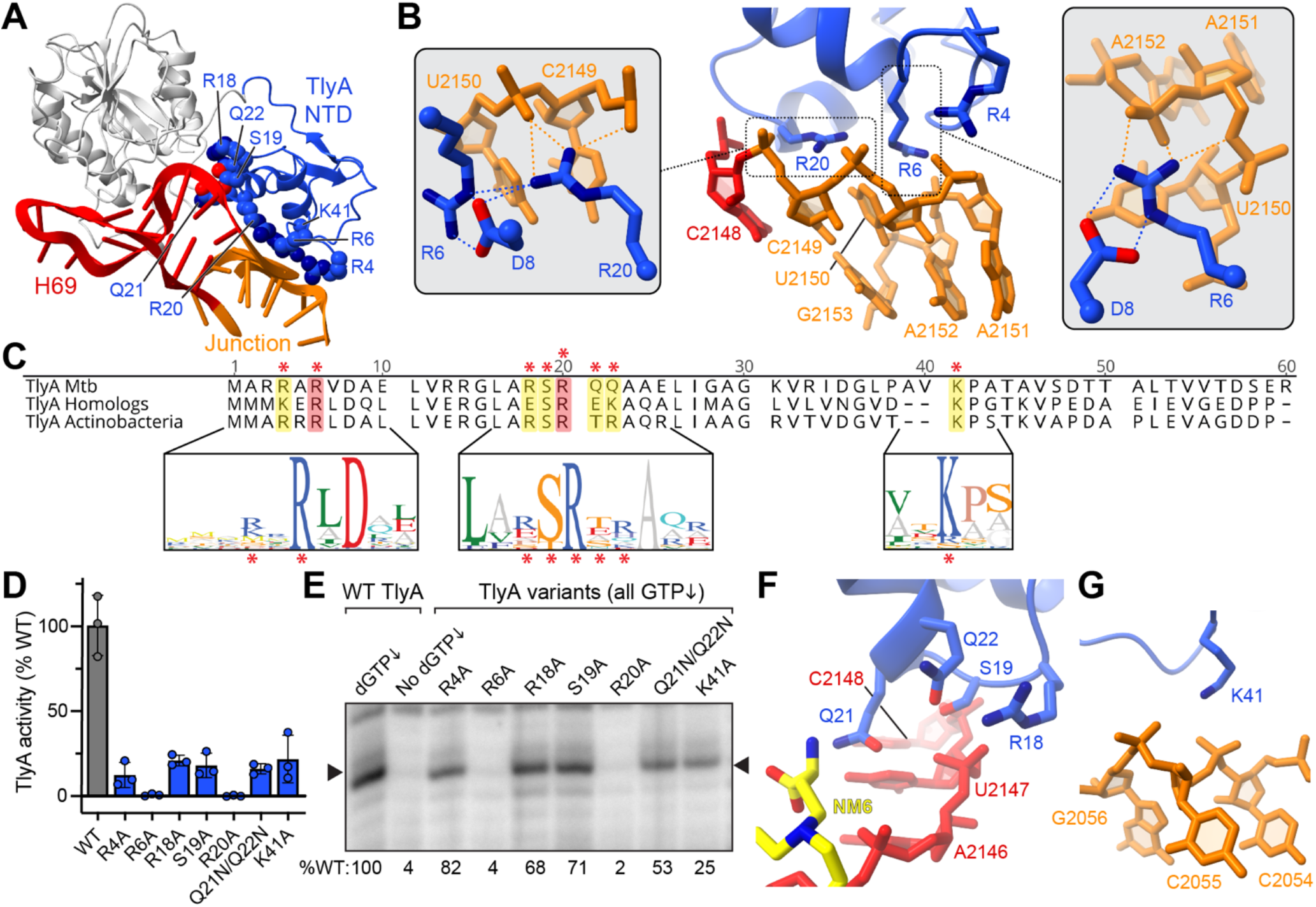
The TlyA NTD recognizes a complex rRNA structure at the base of H69. ***A***, Overview of the TlyA-H69 complex highlighting NTD residues on the TlyA interaction surface for which amino acid substitutions were made. ***B***, Zoomed in view of interactions made by TlyA NTD residues Arg4, Arg6 and Arg20 with nucleotides of the rRNA junction proximal to H69 (orange). Insets: alternate views of Arg6/Arg20 and Arg6 alone with interactions with rRNA and between protein residues indicated with orange and blue dotted lines, respectively. ***C***, Sequence alignment of the *Mtb* TlyA NTD sequence with the consensus sequences for all TlyA homologs and closer homologs from actinobacteria only. Shown below are sequence logo plot representations of sequence conservation for the selected regions among actinobacterial TlyA sequences. The red asterisk denotes sites of amino acid substitutions generated in this work. ***D***, *In vitro* methylation of *Msm* 50S subunit by wild-type (normalized to 100%) and variant TlyA proteins using [^3^H]-SAM. ***E***, Representative gel showing the results of *Msm* 50S subunit methylation by variant TlyA proteins detected via RT using a radiolabeled DNA primer. Stops at methylated C2144 ribose (arrowhead) are only observed under conditions of depleted dGTP (compare first two lanes with wild-type TlyA). Values below the image are the average band intensity relative to wild-type TlyA for at least two independent experiments. Zoomed views of the interactions made by TlyA NTD residues ***F***, Arg18, Ser19, Gln21 and Gln22, and ***G***, Lys41.

To confirm the importance of Arg6 and Arg20 residues in 23S rRNA recognition, individual alanine substitution variants were created, and their proper folding was confirmed using nano-differential scanning fluorimetry (nDSF; **Fig. S8**). Next, enzyme activity was assessed in two complementary methyltransferase activity assays: quantification of 50S subunit ^3^H incorporation following transfer of a [^3^H]-methyl group from radiolabeled SAM cosubstrate ([^3^H]-SAM) and direct visualization of C2144 2’-O-methylation via reverse transcription (RT) primer extension. For the [^3^H]-SAM assay, we first established optimal conditions using wild-type TlyA and then compared these and all other variants in a single time-point assay under conditions corresponding to ~90% completion of 50S subunit methylation for the wild-type enzyme (**Fig. S9**). Consistent with an essential role in specific 50S subunit substrate recognition, individual substitution of either Arg6 or Arg20 completely eliminated methyltransferase activity (**Fig. 3D**). This result was corroborated in the RT assay in which no methylation above background at C2144 was observed for either TlyA R6A or R20A (**Fig. 3E**). In contrast, TlyA R4A exhibited some activity in the [^3^H]-SAM assay and more robust methylation via primer extension, suggesting this residue makes a smaller contribution to 50S subunit binding by TlyA, as previously noted (17). While the reason for the difference between the two assays in R4A variant activity is not immediately obvious, the RT assay is less readily amenable to accurately assessing complete methylation for wild-type TlyA and thus for quantitative comparison with variants.

Four other residues surrounding Arg20 are also positioned to make interactions with H69 nucleotides A2146, U2147 and C2148, as well as the TlyA-bound SAM analog. Arg18 is adjacent to one non-bridging oxygen of the phosphate group of U2147, while Gln21 is located between the second non-bridging oxygen of the same phosphate group, the base O4 atom of U2147, and the terminal carboxyl group of the bound SAM analog (**Fig. 3F; Fig. S6B**). Ser19 is also located between these residues and the phosphate of C2148, with Gln22 in a central location within 3-4 Å of all three other TlyA residues as well as the U2147 phosphate group. As before, these residues were substituted with alanine (R18A and S19A) or as a double change with a more conservative asparagine substitution at both glutamine residues (Q21N/Q22N) and assessed in the two activity assays after confirming their correct folding (**Fig. 3D,E; Fig. S8B,D**). Consistent with their more modest conservation among TlyA homologs compared to Arg6 and Arg20, all three variant proteins were affected by the amino acid substitution but retained some activity in both assays (**Fig. 3C-E; Fig. S7**). These results suggest these residues play supporting, but not individually critical, roles in TlyA substrate recognition

Finally, within the TlyA NTD, the highly conserved Lys41 is positioned to make a single electrostatic interaction with the phosphate group of C2055 which is in a bulge loop at the base of H68, on the strand complementary to the 23S rRNA sequence preceding H69 (**Fig. 3C,G; Fig. S6C, Fig. S7**). Again, a reduction in both activity assays was observed suggesting an important, but not individually critical, role in 50S subunit binding for Lys41 (**Fig. 3D,E**). Thus, these analyses have identified the NTD residues that contribute collectively to 50S subunit interaction (Arg4, Arg18, Ser19, Gln21, Gln22, and Lys41), including two, Arg6 and Arg20, whose coordinated recognition of a complex 23S rRNA structure adjacent to H69 is critical for specific substrate recognition by TlyA.

### TlyA CTD interactions with H69 position the methyltransferase domain for C2144 modification

The TlyA CTD makes extensive contact with the irregular minor groove of H69, from the U2132:A2146 pair near the base to the tip of the helix, with two positive patches on either side of the SAM binding pocket extending the NTD contact surface on the rRNA (**Fig. 2**). Five TlyA residues of moderate to very high conservation are positioned to interact with H69: Arg65, Tyr115, Arg133, Arg137, and Lys189 (**Fig. 4A-D; Fig. S6D-F**). Two additional residues at the C2144 target nucleotide, Phe157 and Ser234, and their role in TlyA activity are described further in the next section. As before, each residue was individually substituted with alanine, and additionally to isoleucine in the case of Tyr115 to specifically probe the requirement for an aromatic side chain at this position. The purified variant proteins were assessed using nDSF (**Fig. S8**) which revealed them to be properly folded with only one potential exception, Y115I, which retained an unfolding temperature (T_i_) similar to the wild-type protein but with an inverted profile (**Fig. S8E**).

**Figure 4.**
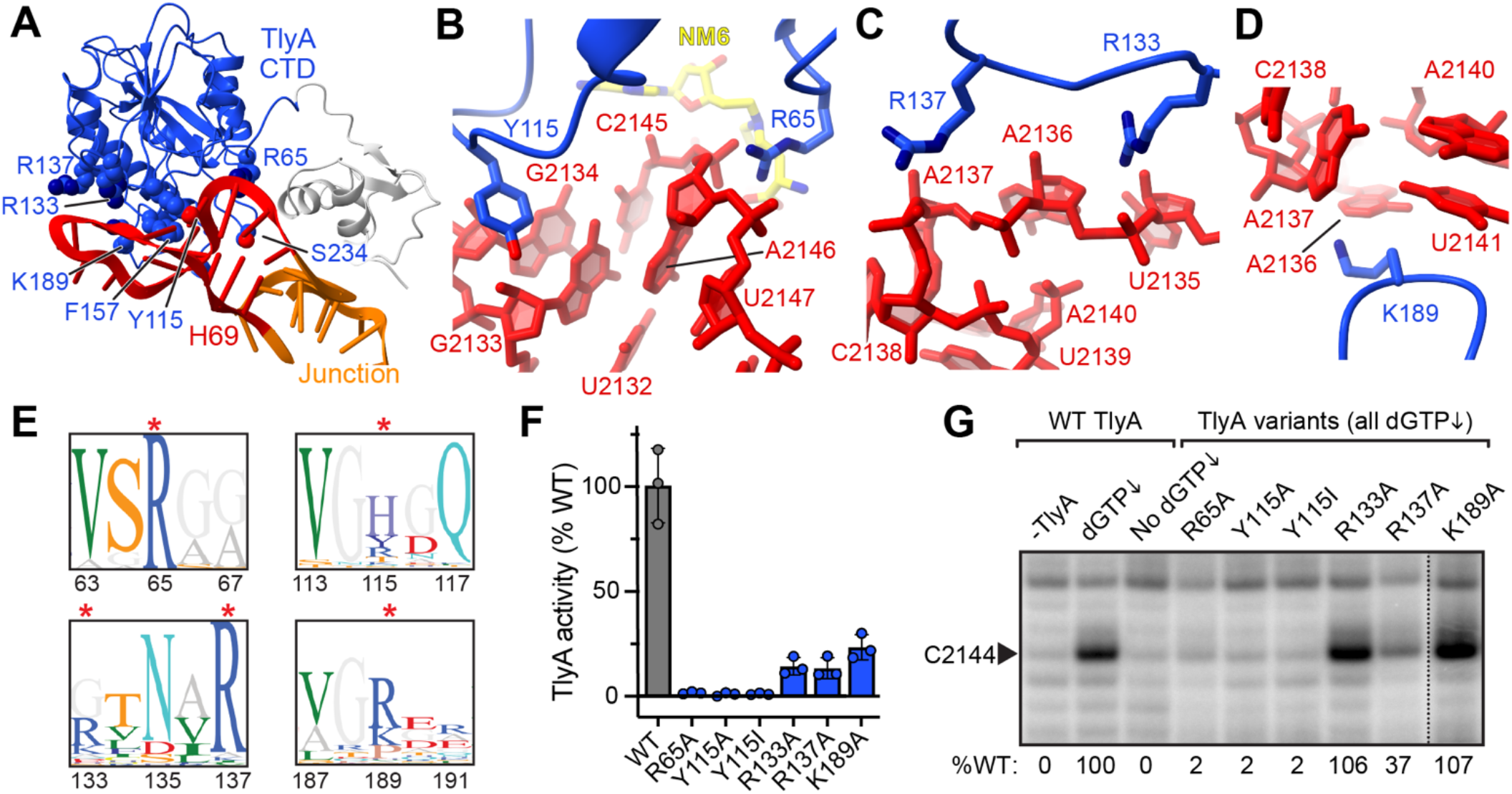
The TlyA CTD interacts with H69 surrounding the modification site. ***A***, Overview of the TlyA-H69 complex highlighting CTD residues on the TlyA interaction surface for which amino acid substitutions were made. Zoomed in views of interactions between H69 (red) and TlyA CTD residues ***B***, Arg65, Tyr115, ***C***, Arg133, Arg137, and ***D***, Lys189. ***E***, Sequence logo plot representations of actinobacterial TlyA sequence conservation for regions surrounding the selected CTD residues. The red asterisk denotes sites of amino acid substitutions made in this work. ***F***, *In vitro* methylation of *Msm* 50S subunit using [^3^H]-SAM for wild-type TlyA and CTD variant proteins. ***G***, Representative gel showing the results of RT analysis of *Msm* 50S subunit methylation by TlyA CTD variants. Values below the image are the average band intensity relative to wild-type TlyA for at least two independent experiments.

Towards the base of H69, Arg65 recognizes the phosphate backbone of nucleotides A2146 and U2147 with the guanidine head group positioned beneath the phosphate of A2146 and within electrostatic interaction distance of a non-bridging oxygen of the U2147 phosphate (**Fig. 4B**). On the opposite stand of H69, Tyr115 extends into the minor groove, contacting the G2134 ribose and G2133 ribose and base edge, with its hydroxyl group within hydrogen bonding distance of the G2133 nucleobase N3 atom (**Fig. 4B**). Positioning of Tyr115 to make these interactions also depends upon a local but significant loop reorganization between the free and 50S subunit-bound forms of TlyA (**Fig. S5B**). Arg65 is universally conserved and substitution with alanine completely ablates activity in both assays (**Fig. 4E-G; Fig. S7**), consistent with a critical role in 50S subunit binding and substrate recognition. Substitution of Tyr115 with either alanine or isoleucine similarly results in fully diminished enzyme activity in both assays (**Fig. 4F-G**). Although Tyr115 is not as highly conserved as Arg65, this position is most commonly aromatic and/ or basic (e.g. tyrosine, histidine or arginine; **Fig. 4E; Fig. S7**), suggesting conservation of interactions like those we observe in the structure is essential in other TlyA homologs.

Three basic TlyA residues (Arg133, Arg137, and Lys189) surround the hairpin loop structure at the tip of H69. Arg133 and Arg137 approach the backbone phosphate groups on the minor groove side of nucleotides U2135/A2136 and A2138, respectively, while on the opposite side of H69, Lys189 is positioned alongside the base edges of A2136 and U2141 (**Fig. 4C,D**). Arg137 is highly conserved among TlyA homologs (85-90%), whereas conservation is more modest at the other two positions, though still most commonly a basic residue (~40-77% Arg/Lys; **Fig. 4E; Fig S7**). Functional analyses of individual alanine substitution variants at these residues revealed a modest impact on TlyA activity with all three comparably reduced in the [^3^H]-SAM assay, but only R137A exhibiting significantly diminished activity in the RT assay (**Fig. 4F,G**). While these results suggest that Arg133, Arg137, and Lys189 contribute to H69 binding by TlyA, this may be accomplished through their collective interactions with the tip of the helix.

Together, our structural insights and functional analyses suggest that the TlyA CTD contains at least two residues critical for H69 binding, Arg65 and Tyr115, and several others that collectively recognize features along the length of H69. Further, these residues lie on a contiguous surface with similarly essential NTD residues (Arg6 and Arg20), suggesting coordinated recognition of distinct features of 23S rRNA underpin specific substrate recognition of the 50S subunit by TlyA.

### TlyA employs a base flipping mechanism to position C2144 for ribose methylation

Binding of TlyA on the 50S subunit precisely positions the opening to the SAM binding pocket and the TlyA active site directly over the target nucleotide C2144 (**Fig 5A**). As noted earlier, use of NM6 in preparing the 50S-TlyA complex also facilitated capture of the enzyme in a post-catalytic state, with C2144 covalently modified on its 2’-OH and the SAM analog still bound in TlyA’s cosubstrate binding pocket. While much of H69 and the adjacent rRNA junction is structurally unaltered upon TlyA binding, suggesting that the enzyme specifically recognizes the mature 50S subunit, we observe significant local deformations around the target nucleotide in our structure. Most strikingly, C2144 fully flips out from H69, with two TlyA residues, Phe157 and Ser234, positioned to stabilize the altered H69 structure (**Fig. 5B-D; Fig. S6G**). Phe157, which is almost universally conserved among TlyA homologs (**Fig. 5C, Fig. S7**), stacks on A2143 and partly fills the space normally occupied by C2144. This interaction appears mechanistically critical as removal of the amino acid side chain (F157A substitution) completely abrogates activity (**Fig. 5D-F**). Further, a F157I substitution, which maintains a bulkier side chain but lacks an aromatic nature that would favor stacking on the RNA base, also renders TlyA completely inactive, suggesting that the π-π stacking of aromatic side chain and nucleobase is specifically critical. In contrast, the observed interaction of Ser234 via its hydroxyl group with the NH_2_ of C2144 is not essential for activity, consistent with the very low level of conservation at this position (**Fig. S7**). As such, TlyA does not appear to require direct identification of the base identity at C2144 for modification (**Fig. 5D-F**). However, we also note that Ser234 is flanked by two universally conserved glycine residues which likely impart the necessary flexibility in this short loop to intimately sequester the flipped base which may allow some level of discrimination among possible RNA bases.

**Figure 5.**
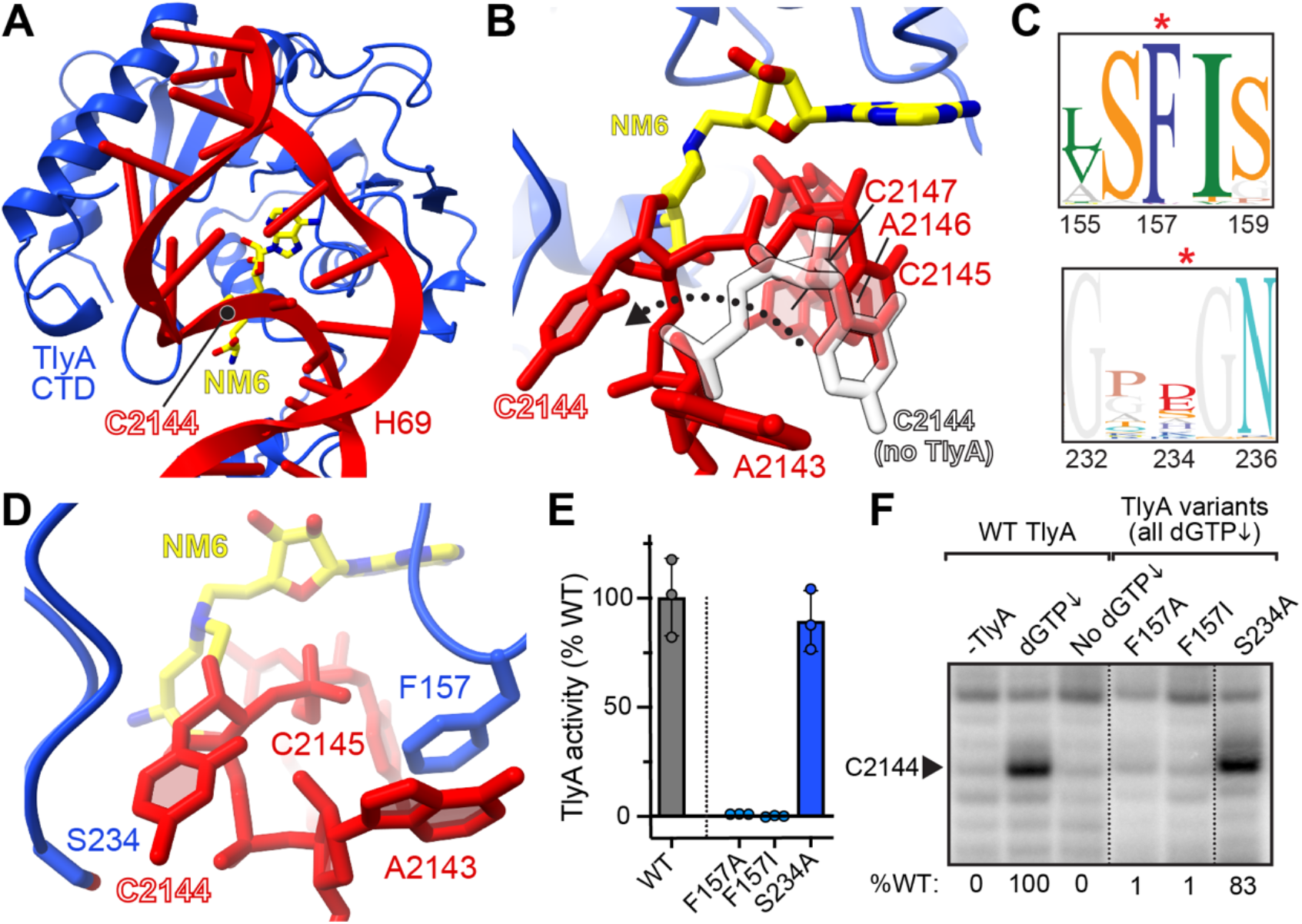
TlyA uses a base flipping mechanism to position C2144 for 2’-OH modification. ***A***, View of TlyA CTD and NM6 cosubstrate positioned over the H69 modification site. ***B***, The NM6-modified C2144 nucleotide is flipped from H69 compared to its original position (white, semi-transparent sticks). ***C***, Sequence logo plot representations of actinobacterial TlyA sequence conservation for regions surrounding the selected CTD residues proximal to the flipped nucleotide. The red asterisk denotes sites of amino acid substitutions generated in this work. ***D***, TlyA CTD residues Phe157 and Ser234 interact with H69 nucleotide C2144 and A2143, respectively. ***E***, *In vitro* methylation of *Msm* 50S subunit using [^3^H]-SAM for wild-type TlyA and indicated CTD variant proteins. ***F***, Representative gel showing the results of RT analysis of *Msm* 50S subunit methylation by these TlyA CTD variants. Values below the image are the average band intensity relative to wild-type TlyA for at least two independent experiments. In *panels E* and *F*, data for wild-type TlyA are the same as in **Fig. 4** (dotted lines denote regions removed from the original images).

### Insights into 30S subunit recognition and impact of TlyA clinical mutations

With the new structural and functional understanding presented thus far on how TlyA specifically recognizes the 50S subunit for C2144 ribose modification, we addressed two key questions on TlyA’s dual substrate specificity and the functional impact of known clinical variants that lead to capreomycin resistance. First, using the collection of TlyA variants already generated, we asked whether the same dependencies on specific NTD and CTD residues also applies to substrate recognition and modification of TlyA’s 30S substrate target site (**Fig. 6A**). Most strikingly among the NTD variants, some activity is retained in both the R6A and R20A variants, with the latter exhibiting around 50% activity compared to the wild-type enzyme. This is in stark contrast to modification of the 50S subunit where both amino acid substitutions fully eliminated TlyA activity (**Fig. 3D**). Additionally, the R4A substitution which had more modestly reduced activity on 50S subunit, resulted in an equal reduction in activity to R6A for 30S subunit modification. The activity of CTD variants on 30S subunit also appears to differ with some activity observed for the R65A, Y115A/I, and F157A/I variants which fully abrogated activity on the 50S subunit. However, as for 50S subunit modification, S234 does not appear to play a critical role in substrate recognition. These results suggest that the same molecular surfaces engage with both 50S and 30S subunits, but the critical dependencies on specific residues are distinct for TlyA’s interaction with its two substrates.

**Figure 6.**
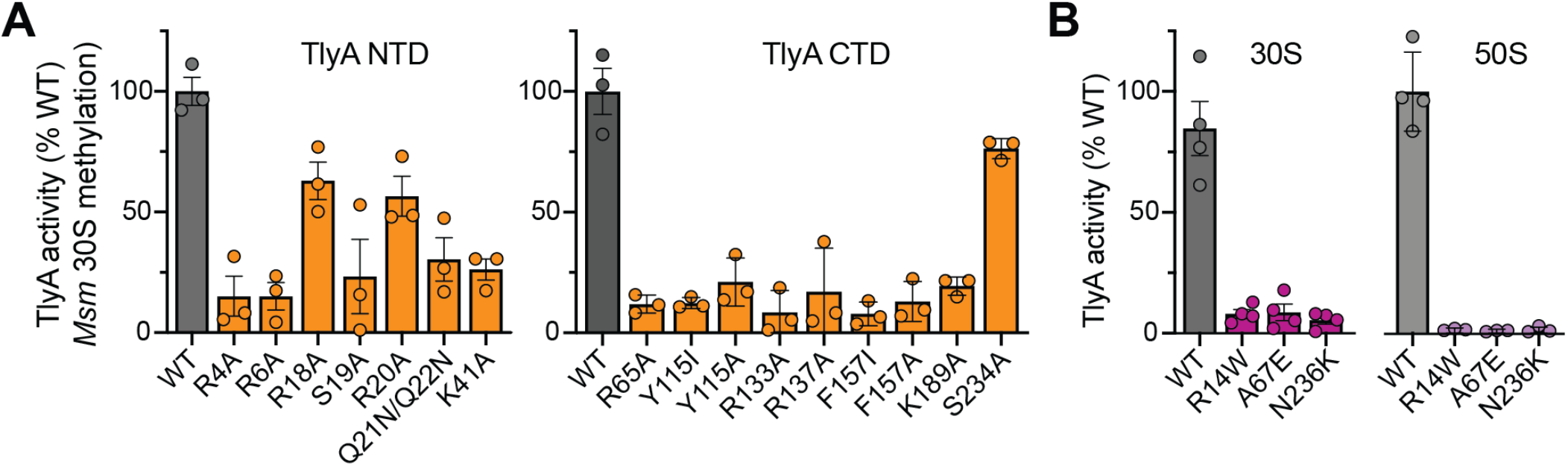
TlyA has distinct residue dependencies for 30S methylation and is inactivated by clinically-identified resistance mutations. ***A***, *In vitro* methylation of *Msm* 30S subunit by wild-type TlyA (normalized to 100%) and the indicated NTD (*left*) and CTD (*right*) variant proteins using [^3^H]-SAM. ***B***, As in *panel A* but for *Msm* 30S (*left*) or 50S (*right*) subunit with the three indicated TlyA variants associated with clinical resistance to capreomycin.

Clinical resistance to capreomycin can arise through 16S rRNA mutation (27, 28) or via amino acid substitutions in TlyA that eliminate its activity and thus incorporation of the rRNA methylation required for optimal capreomycin activity (28–31). To directly test whether these TlyA mutations result in enzymes lacking activity due to structural disruption of key interactions with the ribosome subunits, we created additional TlyA variants corresponding to three *Mtb* mutations associated with capreomycin resistance, R14W, A67E and N236K (28–31). Although all three proteins were expressed and soluble, analysis of their folding using nDSF suggested more significant structural perturbations for R14W and N236K (**Fig. S10**), compared to other NTD and CTD variants designed to test specific interactions as described above. Consistent with their role in clinical resistance to capreomycin, all three TlyA variants were unable to methylate either substrate efficiently, with no detectable activity on the 50S subunit for modification of C2144 (**Fig. 6B**).

## Discussion

Changes in rRNA modification status can have profound effects on ribosome assembly, function, and sensitivity to ribosome-targeting antibiotics (32). rRNA methylations have been identified which either block antibiotic action or are necessary for optimal drug binding and thus antibacterial activity. In bacteria, these rRNA methylations are incorporated by Class I or Class IV methyltransferases (33, 34), with a single enzyme typically responsible for each individual modification. Exceptions to this strict specificity do exist, such as for TlyA which is capable of incorporating cytidine 2’-O-methyl modifications on both the small and large subunit, within two structurally distinct contexts.

Here, we determined the cryo-EM structure of TlyA bound to the 50S ribosomal subunit, revealing the full-length structure of the enzyme and the detailed molecular mechanism of specific recognition of one of its two ribosomal subunit substrates. To our knowledge, this structure represents the first example of an rRNA 2’-O methyltransferase bound to a bacterial ribosomal subunit. These studies also identified an essential 23S rRNA interaction surface that spans both the NTD and CTD of TlyA and contains a set of residues critical for 50S subunit substrate binding and 2’-O-methylation of C2144. TlyA accomplishes specific 50S subunit recognition via essential interactions of its NTD with a unique tertiary structure at the base of H69 and of the CTD with H69, which position the bound methyl group donor SAM over the target nucleotide. Finally, the nucleobase of C2144 is flipped out of the H69 helical stack, in a conformation stabilized by TlyA Phe157 stacking on the adjacent A2143, placing the 2’-OH of the ribose of C2144 adjacent to SAM and the catalytically important residues of TlyA (22).

The TlyA NTD adopts an S4 ribosomal protein fold that makes critical interactions, primarily by the highly conserved TlyA residues Arg6 and Arg20, with the complex 23S rRNA structure of the junction at the base of H69. In the original report of the S4 structure from *Bacillus stearothermophilus*, the corresponding S4 residues (Arg92 and Arg106 in the *Msm* S4 domain 2; **Fig. S5F,G**) were among several highly conserved basic or aromatic residues proposed to form an extensive rRNA binding surface (35). However, S4 Arg92 and Arg106 do not make extensive interactions with 16S rRNA in the *Msm* 70S ribosome structure and instead these residues appear to play important roles in S4 interdomain interactions (25). In contrast, two other S4-domain containing proteins involved in ribosomal quality control, YabO from *Bacillus subtilis* and the human mitochondrial MTRES1 (36–38), recognize the complex structure at the base of H69 in a manner conserved with TlyA (**Fig. S5H-J**). In particular, the residues corresponding to TlyA Arg6 and Arg20 in the first two α-helices of the YabO and MTRES1 S4 protein folds make essentially the same interactions with the conserved rRNA tertiary structure (**Fig. SJ**). Although MTRES1 appears a little more divergent in sequence, the action of YabO Arg2 (TlyA Arg4) and Arg16 (TlyA Arg20) may also be supported by the conserved Asp4 (TlyA Asp8). Thus, the S4 protein domain thus appears to be a modular unit which has been adopted by diverse proteins for 50S recognition at the base of H69.

Our structure revealed C2144 to be flipped out of H69 in a post-catalytic state captured by use of the SAM analog. Base flipping is a common strategy used by DNA modifying or repair enzymes and has been observed or proposed for other rRNA modifying enzymes (39–42). However, given its relative accessibility in the RNA minor groove and as a common component of all RNA nucleotides, whether this molecular strategy would be employed for ribose 2’-OH methylation was previously less clear. Our structure suggests that base flipping may be a common mechanistic feature regardless of RNA methylation target site. Such a strategy would provide an opportunity to probe the base identity for specific target site recognition and could optimally adjust the 2’-OH geometry for methylation. Ribose methylation can influence sugar pucker and base flipping (43), and may be an important element of the modification process itself.

Comparisons to the well characterized 16S rRNA (m^1^A1408; *E. coli* numbering) aminoglycoside-resistance Class I methyltransferases, such as NpmA (42), are also particularly intriguing. NpmA also uses a base-flipping mechanism despite the N1 atom being relatively exposed on the helix 44 surface. Like TlyA, NpmA relies heavily on recognition of complex rRNA structure, distant from the site of modification, to accomplish specific binding to its substrate (42). Further, a single direct base edge contact is made by NpmA to A1408, but the absolute importance of this interaction is unclear given that NpmA retains partial activity against ribosomes with a G1408 nucleotide (44). Similarly, TlyA contacts the nucleobase of C2144 via Ser234 located in the loop linking the sixth and seventh β-strands (β6/7 linker) of its Class I methyltransferase core fold, a region commonly associated with substrate recognition by these enzymes (34, 45). As for NpmA, this specific base contact does not appear critical for TlyA activity based on our functional analyses. However, as noted earlier, the high conservation of the surrounding sequence suggests the TlyA β6/7 linker structure may nonetheless be important for forming the pocket shielding C2144 from exposure to solvent in its flipped conformation. One distinction between TlyA and NpmA appears to be how the flipped conformation is stabilized. In NpmA, a basic residue (Arg207) stabilizes a local distortion of the h44 backbone, and the flipped A1408 is stabilized by stacking between two conserved tryptophan residues. Additionally, the vacated space within helix 44 is left unoccupied and NpmA does not contact or stabilize bases on the complementary strand. In contrast, TlyA uses the conserved Phe157 to occupy the space vacated by flipping of C2144 via stacking on the adjacent A2143 nucleobase. As such, the mechanism used by TlyA is more akin to DNA methyltransferases which replace DNA base pairing and stacking interactions, normally made by the flipped base, with protein-DNA contacts to the base left unpaired within the DNA double helix (46).

The similarities and distinctions between TlyA and NpmA may also be significant for TlyA’s mechanism of recognition of its other target nucleotide in the 30S subunit, C1392 (C1409 in *E. coli*), which immediately follows A1408 in the 16S rRNA. Our speculation is that TlyA may exploit the same complex 16S rRNA tertiary surface used by NpmA and related enzymes, as well as the m^7^G1405 aminoglycoside-resistance methyltransferases (47). Our analysis of 30S methylation by the NTD and CTD variants of TlyA suggest that the same surfaces containing these altered residues are also broadly engaged in recognition of the 30S subunit. However, some important differences in dependencies on specific key residues for substrate interaction are apparent: whereas Arg20 is essential for 50S modification on C2144, alteration of this residue only minimally impacts 30S methylation. In contrast, 16S rRNA C1392 modification appears to depend more on additional residues at the very N-terminus (e.g. Arg4). This finding is also consistent with insights gleaned from the existence of two TlyA subtypes, TlyA^I^ and TlyA^II^, of which only the longer TlyA^II^ possesses dual substrate specificity and is able to modify the 30S subunit. TlyA^I^ enzymes lack a short sequence at their N-terminus (containing Arg4) and an entire α-helix that follows the seventh core β-strand in TlyA^II^. Thus, consistent with our functional assays and previous alterations of Arg3 and/ or Arg4 (17), critical elements of 30S subunit recognition appear to reside in these regions. Precisely how TlyA adapts to the two structurally distinct target sites remains to be fully elucidated, but we have also previously proposed that structural plasticity in the short interdomain linker in TlyA may be a mechanism by which the enzyme could accomplish this (21). In further support of this idea, a known clinical capreomycin resistance mutation resulting in an A67E substitution in TlyA (28), which we found to inactivate the enzyme, would likely disrupt the hydrophobic binding pocket which linker residue Trp62 occupies in the 50S subunit-bound TlyA structure. This change, in turn, could prevent SAM binding or correct NTD/ CTD association or interdomain communication. The present work thus reveals common requirements in TlyA for modification of both its substrates, but with some key differences in the residues most critical for individual subunit recognition, and adds support to a mechanism by which TlyA might structurally adapt to these distinct interaction surfaces. However, fully defining the basis of TlyA’s dual substrate specificity will require corresponding detailed structural studies of TlyA and its 30S subunit substrate.

Of the two modifications incorporated by TlyA, C2144 methylation most strongly influences the binding of capreomycin (approximately 20 Å away) by a long-range mechanism that is not currently well defined. Comparison of H69 in multiple ribosome structures available in the PDB with and without C2144 modification reveals a small but consistent difference at the tip H69: in unmodified ribosomes, the loop formed by nucleotides A2137, C2138, and U2139 makes a tighter turn than in modified ribosomes. Our structure now offers a third comparison, with a bulkier modification incorporated but with TlyA still also bound, in which H69 is observed in an structural state between those of ribosomes with unmodified and modified C2144. In the conformation of other unmodified bacterial ribosomes, when H69 is more tightly bent, the bases of A2137 and C2138 are more distant from the capreomycin binding site on the 30S subunit. These observations suggest that modification of C2144 alters the structure of H69 in a manner that changes the position of nucleotides A2137 and C2138, promoting the direct interactions they make with capreomycin.

In addition to the A67E substitution in TlyA noted above, our work offers insight into how capreomycin resistance arises clinically through two other mutations in the gene encoding TlyA. In the TlyA NTD, the mutation resulting in an R14W substitution (28) likely disrupts TlyA NTD folding and its essential contribution to substrate recognition on the 50S subunit. Although it does not directly contact 23S rRNA, Arg14 is positioned directly above Arg6 and Arg20, and interacts with the TlyA backbone at Thr50/Ala51 which are part of a loop that wraps closely around the arginine side chain. Thus, structural changes to accommodate the bulkier tryptophan side chain would disrupt the critical interactions with the rRNA made by Arg6 and Arg20. Another common mutation found in resistant *Mtb* results is a N236K substitution (30) which our structure suggests could impact TlyA activity in several ways. Gln236 immediately follows the β6/7 linker (sequence ^232^GPSG^235^) which surrounds the flipped C2144 base. Additionally, this substitution places a lysine residue close to residue Glu238 which has been proposed to play an important role in catalysis (22).

In summary, the present work has revealed the full-length structure of the *Mtb* methyltransferase TlyA and defined the molecular basis for specific recognition of its 50S subunit substrate. While future structural and biochemical studies with the 30S subunit will be necessary for a full understanding of TlyA’s dual substrate specificity, these studies have deepened our understanding of rRNA methyltransferase action. In particular, recognition of unusual rRNA structures distant from the site of modification and base flipping both emerge as general themes in substrate molecular recognition for these enzymes.

## Materials and Methods

### TlyA protein expression, purification and site-directed mutagenesis

An *E. coli* codon-optimized sequence encoding *Mtb* (strain ATCC 25618/H37Rv) TlyA was obtained via chemical synthesis (GeneArt) and subcloned into a pET44a(+) plasmid (pET44-TlyA), as previously described (21). This construct produces TlyA with an N-terminal hexahistidine tag. The TlyA-encoding plasmid was used to transform *E. coli* BL21 (DE3) and cultures were grown at 37 °C in Terrific Broth containing 100 μg/mL ampicillin. At mid-log phase (~0.4-0.6 OD_600_) protein expression was induced with 0.5 mM isopropyl β-D-1-thiogalactopyranoside, and growth continued for an additional 3.5 hours. Following harvest via low-speed centrifugation (4,000 × *g*) for 10 minutes at 4 °C, the cells were resuspended in lysis buffer (50 mM NaH_2_PO_4_, pH 8.0, 300 mM NaCl, and 10 mM imidazole containing an EDTA-free SIGMA*FAST*™ Protease Inhibitor Cocktail Tablet) and lysed by sonication (Misonix Sonicator 3000 with microtip: 15 minutes total sonication time, 0.9 s on, 0.6 s off, output level 5.5). Cell lysates were cleared by centrifugation (21,000 × *g*) at 4 °C for 40 minutes and filtered before purification of TlyA by sequential Ni^2+^-affinity (Cytiva HisTrap™ FF crude 1mL or manual His-column using Millipore Ni-NTA His-Bind® Resin) and gel filtration (Cytiva HiLoad™ 16/600 Superdex™ 75) chromatographies on an ÄKTA Purifier 10 system. TlyA variants with single or double amino acid substitutions were created using megaprimer whole-plasmid PCR (48) in pET44-TlyA, and expressed and purified by Ni^2+^-affinity chromatography as described above for wild-type TlyA. Protein folding and quality control was accomplished using nDSF on a Tycho NT.6 (NanoTemper) which monitors protein thermal unfolding using intrinsic fluorescence at 330 and 350 nm. The unfolding profile and inflection temperature (T_i_) of each TlyA variant was determined using the instrument software for comparison to that of wild-type TlyA.

### Isolation of Msm 50S and 30S subunits

*Msm* 50S subunits with unmethylated C2144 were isolated from a strain lacking TlyA activity (LR222 C101A) using following established procedures (21, 49). A small culture of Middlebrook 7H9 liquid medium was inoculated with a single colony of *Msm* LR222 C101A and grown overnight at 37 °C with shaking (100 rpm). Fresh Middlebrook 7H9 medium (0.5-2 L) was inoculated with the overnight culture (1/100 dilution) and the cultures grown for 72 hours at 37 °C with shaking (100 rpm). Cells were harvested by low-speed centrifugation (4,000 x *g*) for 10 minutes at 4 °C and washed (500 mL per L culture) twice with a solution of 10 mM HEPES/KOH (pH 7.6), 10 mM MgCl_2_, 1 M NH_4_Cl, and 6 mM 2-mercaptoethanol, and once with the same solution but with only 0.1 M NH_4_Cl. The cells were then resuspended in the same final buffer and lysed using three passages through a French Press. After addition of DNAse I (10U/ ml lysate), the lysate was cleared by centrifuging for 10 and 30 minutes (at 17,300 and 26,900 x *g*, respectively), and the resulting supernatant centrifuged at high speed for 18 hours (277,200 x *g*) to pellet ribosomes. The 70S pellet was resuspended and dialyzed against a solution containing 10 mM HEPES/KOH (pH 7.6), 0.3 mM MgCl_2_, 100 mM NH_4_Cl, and 6 mM 2-mercaptoethanol to split the ribosome subunits. 30S and 50S subunits were then separated by centrifugation (90,200 x *g*) on a 10-30% sucrose gradient for 18 hours at 4 °C. The resulting gradient was fractionated using an ÄKTA Purifier 10 system to collect isolated 50S and 30S subunits. Subunits were stabilized by addition of MgCl_2_ to 10 mM and the solution centrifuged (300,750 x *g*) for 18 hours. The resulting individual subunit pellets were resuspended in a solution of 10 mM HEPES/KOH (pH 7.6), 10 mM MgCl_2_, 100 mM NH_4_Cl, and 6 mM 2-mercaptoethanol, flash frozen, and stored at −80°C.

### Cryo-EM sample preparation, data collection and structure determination

SAM analog NM6 (5’-(diaminobutyric acid)-N-iodoethyl-5’-deoxyadenosine ammoniumhydrochloride) was prepared essentially as previously described (24) and purified by semi-preparative reverse-phase HPLC. A 3.0 μL mixture of purified *Mtb* TlyA, *Msm* 50S subunit, and NM6 (at 0.5 μM, 5 μM, and 10 μM respectively) was applied to glow-discharged Quantifoil Cu R1.2/1.3 300 mesh grids. Grids were blotted at room temperature for 3.0-3.3 s at >90% humidity and frozen in liquid ethane using a CP3 plunger (Gatan). Cryo-EM data (3364 micrographs) were recorded as movies with defocus range of −0.8 to −2.2 μm at 81,000x magnification (1.0691 Å/pixel) on a Titan Krios 300 kV (TEM) with Gatan K3 direct electron detector at the National Center for CryoEM Access and Training (NCCAT). The dose per frame was 1.25 e/Å/frame (total dose of 50.79 e^−^/Å^2^) over a total exposure of 2 s divided over 40 frames (50 ms per frame).

Following the workflow outlined in **Fig. S2**, image alignment and dose-weighting were performed using Motioncor2 (50) and RELION-3.0/3.1 (51) was used for subsequent data processing. The contrast transfer function was estimated using the program Gctf (52). To guide automatic picking, 1094 particles were manually picked and then classified into 2D classes. Automatic picking then selected 1,016,454 particles which were extracted with a box size of 280 Å. Multiple rounds of 2D classifications were performed to remove non-ribosomal particles before 3D refinement using a 60 Å low-pass filtered reference map of the *E. coli* 50S subunit (EMD-3133). Iterative rounds of CTF refinement, 3D refinement, and 3D classification were performed resulting in a 3.05 Å post-processed map (**Fig. S2B**, *center*). Analysis of the angular distribution of particles used to generate the map indicated some orientation preference in the data set, but there was good coverage of views containing TlyA (**Fig. S2C**).

Prior to the final post-processing of the complete 50S-TlyA map, multibody refinement was also performed on the remaining particles with separate masks corresponding to TlyA/H69 and the remainder of the 50S subunit, resulting in 3.89 Å and 3.02 Å resolution maps, respectively (**Fig. S2**, *right*). The final three maps (complete 50S-TlyA, TlyA/H69, and 50S subunit alone) were then post-processed using Relion resulting in final 3.05 Å, 3.61 Å, and 2.99 Å maps, respectively, based on gold-standard refinement Fourier Shell Correlation (0.143 cutoff) (**Fig. S2, S3**). Local resolution maps were also generated using ResMap 1.1.4 (53).

All three final maps were used for model building and refinement. The 50S subunit model was created by docking an existing *M. smegmatis* 50S subunit structure (PDB code 5O60), after *de novo* modeling of the NM6-modified C2144, into the 50S-TlyA map and using Coot (version 0.9-pre EL, ccpem) (26) and Phenix (version 1.19.2-4158-000) (54, 55). The TlyA model was generated using a TlyA CTD crystal structure (PDB code 5KYG) appended with a homology modeled NTD (21, 22). As initial docking of our hybrid model (21) did not give a satisfactory fit of the NTD into its portion of the map, this ~60 residue domain was manually re-built. The resulting complete full-length TlyA structure was then used as a search query in the Dali Protein Structure Comparison server (56). This search returned the unpublished structure of a putative hemolysin from *Streptococcus thermophilus* (PDB code 3HP7) as the closest structural homolog which was used to guide further improvement of our TlyA NTD model in regions of the map that were less well resolved. The model was subsequently split into separate TlyA-H69 and remaining 50S subunit models and each separately real-space refined in Phenix (57, 58) using their respective multibody maps (using rigid-body and then to non-rigid body refinement). Finally, the refined models were recombined (without refinement) to create a final complete model of the 50S-TlyA complex and validated using Phenix (54, 55). Complete parameters for data collection and processing, and model building, refinement and validation are summarized in **Table 1**.

**Table 1.** Cryo-EM data collection, refinement and model validation for the 50S-TlyA complex.

|  |  |  |  |
| --- | --- | --- | --- |
| **Deposition** |  |  |  |
| Coordinates (RCSB PDB) | 7S0S |  |  |
| Map (EMDB) | EMD-24792 |  |  |
| **Data Collection/ Processing** |  |  |  |
| Microscope | TFS Titan Krios |  |  |
| Camera | Gatan K3 |  |  |
| Voltage, kV | 300 |  |  |
| Magnification | 81,000x |  |  |
| Electron exposure, e<sup>-</sup>/Å<sup>2</sup> | 50.79 |  |  |
| Defocus range | -0.8 to -2.2 |  |  |
| Pixel size, Å | 1.069 |  |  |
| Symmetry | C1 |  |  |
| No. particles |  |  |  |
| Initial | 1,016,454 | **Multibody Refinement** |  |
| Final | 129,011 | **50S subunit** | **TlyA/H69** |
| Map resolution (FSC 0.143), Å | 3.05 | 2.99 | 3.61 |
| **Refinement and Model** |  |  |  |
| Model resolution (FSC 0.143), Å | 3.0 | 3.04 | 3.66 |
| CC<sub>mask</sub> | 0.66 | 0.84 | 0.69 |
| Map sharpening, Å<sup>2</sup> |  |  |  |
| 50S-TlyA | 83.1 |  |  |
| TlyA-H69 (multibody) | 109.5 |  |  |
| 50S subunit alone (multibody) | 72.9 |  |  |
| Non-hydrogen atoms |  |  |  |
| Protein residues | 3,946 | 3682 | 268 |
| RNA residues | 3,236 | 3213 | 26 |
| Ligand/ modified nucleotide | 407 | 0 | 1 |
| B factors (min/max/mean), Å<sup>2</sup> |  |  |  |
| Protein | 0.21/ 172.4/ 17.1 | 0.21/ 34.9/ 10.4 | 50.0/ 172.4/ 114.4 |
| RNA | 0.05/ 190.2/ 19.7 | 0.05/ 88.6/ 19.2 | 23.1/ 190.2/ 86.7 |
| Ligand/ modified nucleotide | 0.78/ 109.5/ 17.3 | 0.78/ 43.0/ 6.20 | 109.5/ 109.5/ 109.5 |
| RMS deviations |  |  |  |
| Bond lengths, Å | 0.008 | 0.007 | 0.003 |
| Bond angles, ° | 0.915 | 0.808 | 0.760 |
| **Validation** |  |  |  |
| MolProbity score | 1.99 | 1.90 | 2.37 |
| Clashscore | 9.68 | 9.81 | 18.99 |
| Poor rotamers, % | 1.21 | 0.03 | 0.00 |
| Protein |  |  |  |
| Ramachandran plot |  |  |  |
| Favored, % | 93.79% | 94.27 | 88.35 |
| Allowed, % | 6.21% | 5.73 | 10.90 |
| Disallowed, % | 0% | 0.00 | 0.75 |
| RNA |  |  |  |
| Pucker outliers, % | 0% |  |  |
| Bond outliers, % | 0% |  |  |
| Angle outliers, % | 0.1% |  |  |

### RT analysis of 23S rRNA methylation

Extent of methylation of the 50S subunit by wild-type TlyA and mutants was determined using a RT assay. Wild-type or variant *Mtb* TlyA (66 pmol, 2 μM) was incubated for 20 minutes at 37 °C with *Msm* 50S subunit (33 pmol, 1 μM) in the presence of SAM (2.1 μM) in 10 mM HEPES-KOH (pH 7.5), 10 mM MgCl_2_, 50 mM NH_4_Cl, and 5 mM 2-mercaptoethanol. The reaction was terminated by phenol/chloroform extraction and the modified rRNA collected by ethanol precipitation. The rRNA modification at C2144 was assessed using an RT primer-extension reaction with a ^32^P-labeled DNA primer complementary to 23S nucleotides 2188-2204. Modification of the 2’-O was observed only under conditions of complementary (dGTP) depletion (i.e. reactions with 75 μM dATP, dUTP, dCTP; and 0.5 μM dGTP). Controls with no TlyA or no dGTP depletion (i.e. 75 μM dGTP) showed no RT stops corresponding to C2144 ribose modification. Extension products were run on a denaturing (50% urea) 8.6% PAGE sequencing-style gel for 2 hours at 55 W and 50 °C. Gels were dried and then imaged using a phosphor storage screen and Typhoon Trio Variable Mode Imager System (GE Healthcare). Extent of modification was estimated by band intensity comparison using ImageQuant TL 1D Version 7.0.

### Methyltransferase activity assays using [^3^H]-SAM

Quantitative extent of methylation of the 50S subunit by wild-type and variant TlyA was determined using a filter-based enzyme assay with ^3^H-SAM. To establish optimal conditions for comparison to variant proteins, a time-course experiment was performed with wild-type TlyA. TlyA (final concentration 0.76 μM), *Msm* 50S subunits (final concentration 0.38 μM), and ^3^H-SAM (final concentration 0.8 μM) were added to “Buffer G” (5 mM HEPES-KOH (pH 7.5), 50 mM KCl, 10 mM NH_4_Cl, 10 mM MgOAc, and 6 mM 2-mercaptoethanol) to a total reaction volume of 90 μL. The reaction was incubated for 60 minutes at 37 °C, with 10 μL aliquots (3.8 pmol 50S subunit) removed and quenched in 140 μL 5% trichloroacetic acid at 0, 1, 2, 5, 10, 20, 40, and 60 minutes. The reaction was then applied to a glass microfiber filter and 50S subunit methylation quantified using scintillation counting of ^3^H retained on the filter. A 20-minute time point was subsequently selected for comparison of wild-type and variant TlyA proteins using the assay performed essentially otherwise as described above. Assays using 30S subunit were performed using the same procedures, but with single timepoint measurements taken at 60 minutes as activity was observed to be weaker for this substrate (**Fig. S9B**), as previously noted (17).

### Phylogenetic analysis of the TlyA protein family and residue conservation

TlyA homologs were retrieved from InterPro (IPR004538) with conserved sequence feature annotated for Hemolysin A/ rRNA methyltransferase TlyA family. Sequence redundancy was reduced in UniProt using the precalculated UniRef sequence clusters with a cutoff of 50% sequence identity. A total of 223 representative sequences were aligned using Clustal Omega and an unrooted neighbor joining phylogenetic tree was constructed using MEGA X (59) with evolutionary distances computed using the JTT matrix-based method (60). The rate variation among sites was modeled with a gamma distribution (shape parameter = 1) and the residue propensities were calculated using Geneious.

## Acknowledgements

We thank Dr. James Posey and colleagues at the Centers for Disease Control and Prevention, Atlanta GA for providing *M. smegmatis* strain LR222 C101A, and Drs. Puneet Juneja and Ricardo Guerrero-Ferreira for assistance with EM data collection and processing. This work was supported by the National Institutes of Health awards R01-AI088025 (GLC and CMD), T32-AI106699 (ZTL and PS), and T32-GM008602 (PS), and the Burroughs Wellcome Fund Investigator in the Pathogenesis of Infectious Disease award (CMD). This study was supported by the Robert P. Apkarian Integrated Electron Microscopy Core (IEMC) at Emory University, which is subsidized by the Emory School of Medicine and Emory College of Arts and Sciences. Some of this work was performed at the National Center for CryoEM Access and Training (NCCAT) and the Simons Electron Microscopy Center located at the New York Structural Biology Center, supported by the NIH Common Fund Transformative High Resolution Cryo-Electron Microscopy program (U24 GM129539), and by grants from the Simons Foundation (SF349247) and NY State Assembly.

