## Supplementary material for "50S subunit recognition and modification by the *Mycobacterium tuberculosis* ribosomal RNA methyltransferase TlyA": SI

**This PDF file includes:**

Figures S1 to 10

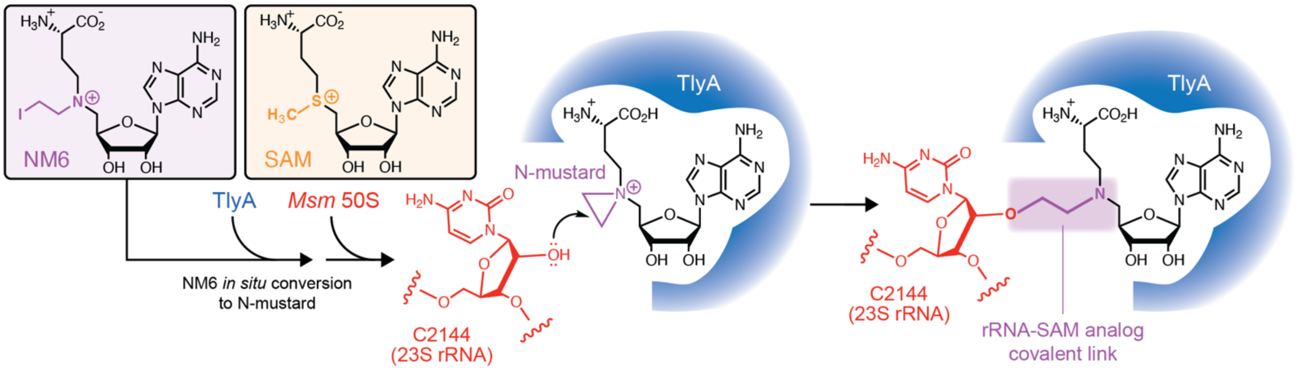


**Fig. S1**. **Stabilization of the 50S-TlyA complex in a post-catalytic state using a SAM analog (NM6).** N-mustard 6 (NM6; *left box*) is an analog of S-adenosyl-L-methionine (SAM; *right box*) that TlyA can use as a cosubstrate for modification of 23S rRNA nucleotide C2144. In this enzymatic reaction, NM6 becomes covalently attached to the ribose 2' position of C2144. The 50S-TlyA complex is thus stabilized in a state immediately following catalysis by virtue of TlyA's affinity for both 50S subunit and the cosubstrate analog covalently attached to C2144.

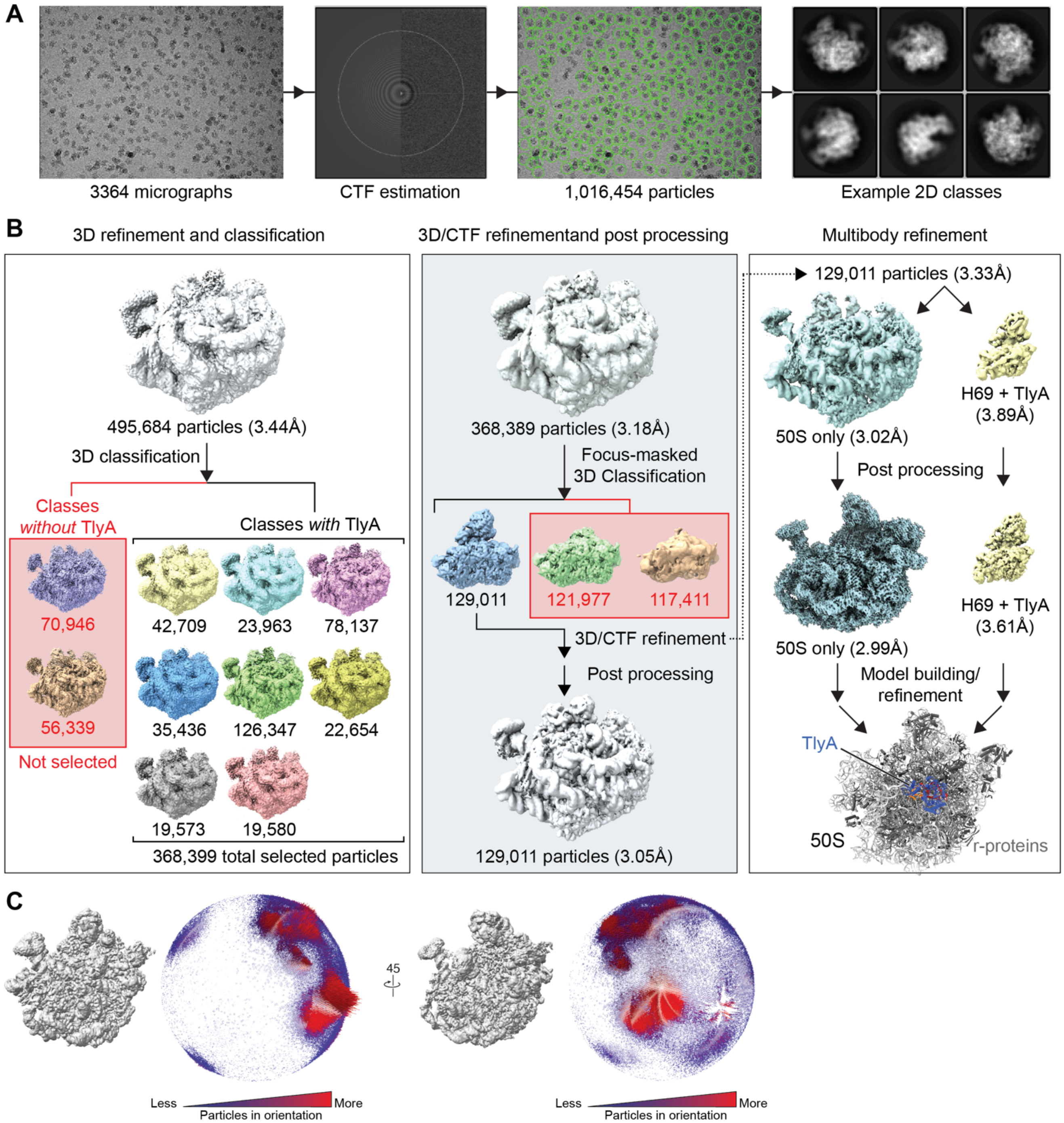


**Fig. S2**. **Workflow for cryo-EM structure determination.** ***A***, Example micrograph, CTF estimation, particle picking, and final 2D classes. ***B***, The 3D refinement workflow leading to the final 50S-TlyA structure. Following initial 3D refinement from 2D classifications (*upper left*), a 3D classification was used to separate classes lacking TlyA (*lower left*). Following a series of subsequent refinement steps (*upper center*), a focus-masked 3D classification was performed to select only high-resolution map for TlyA and the corresponding particles used in a final refinement and post-processing to produce a final map for the 50S-TlyA complex (*lower center*). In addition, using the same map prior to post-processing, a multibody refinement (separating TlyA/ H69 and the remainder of the 50S) was performed to further boost the map resolution in the region of interest (*upper right*). The resulting maps were post-processed, and used for model building and individual refinement in Phenix, before models were combined to generate the final structure (*lower right*). ***C***, Angular distribution plots of particles contributing to the final map. The final map following focus-masked 3D classification (129,011 particles) and corresponding angular distribution plot have the same orientation in both views; the left view is the same as that shown in other figures. The height (low to high) and color (blue to red) of the cylinder bars is proportional to the number of particles in those views.

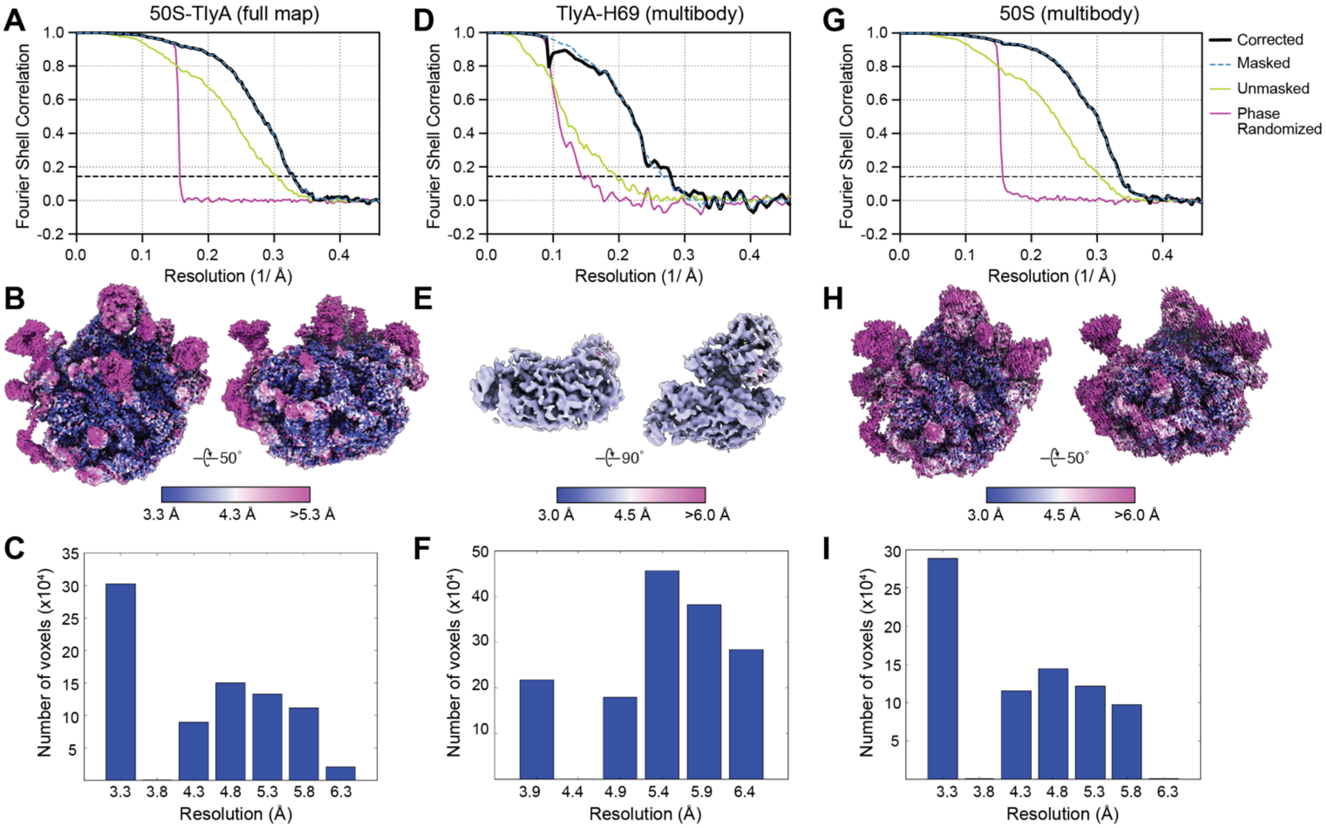


**Fig. S3. Cryo-EM map resolution analysis.** ***A***, Fourier Shell Correlation (FSC) curves for the full 50S-TlyA map: corrected (black), masked data (blue dashed line; under black line where not visible), unmasked data (green), and phase randomized data (purple). Map resolution was calculated from the 0.143 FSC intercept of the corrected data. ***B***, The local-resolution map of the 50S-TlyA complex with resolution indicated as noted in the scale bar. ***C***, Histogram of resolution distribution across the full 50S-TlyA map. Resolution is grouped in 0.5 Å intervals. ***D-F***, as *panels A-C* but for TlyA-H69 multibody refinement. ***G-I***, as *panels A-C* but for 50S-only multibody refinement.

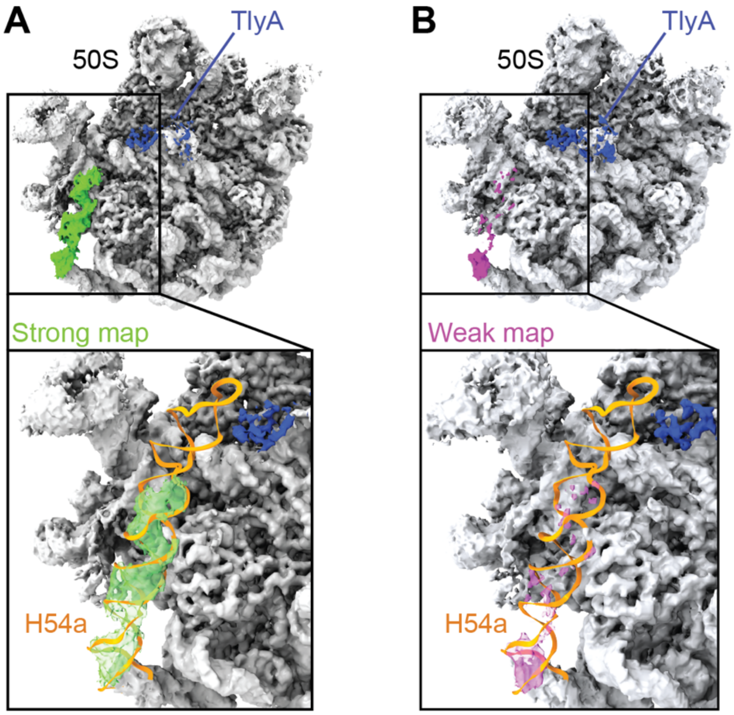


**Fig. S4. Map corresponding to H54a differs among 3D classes.** ***A*,** Map corresponding to 3D reconstruction with H54a (the Handle; green map) visible on the subunit interface surface of the 50S subunit (grey), placing it close to the TlyA NTD (blue map). ***B*,** 3D reconstruction with a disordered H54a and weak map features in the same region (magenta map). In both panels, the final model of H54a (orange) is shown for context.

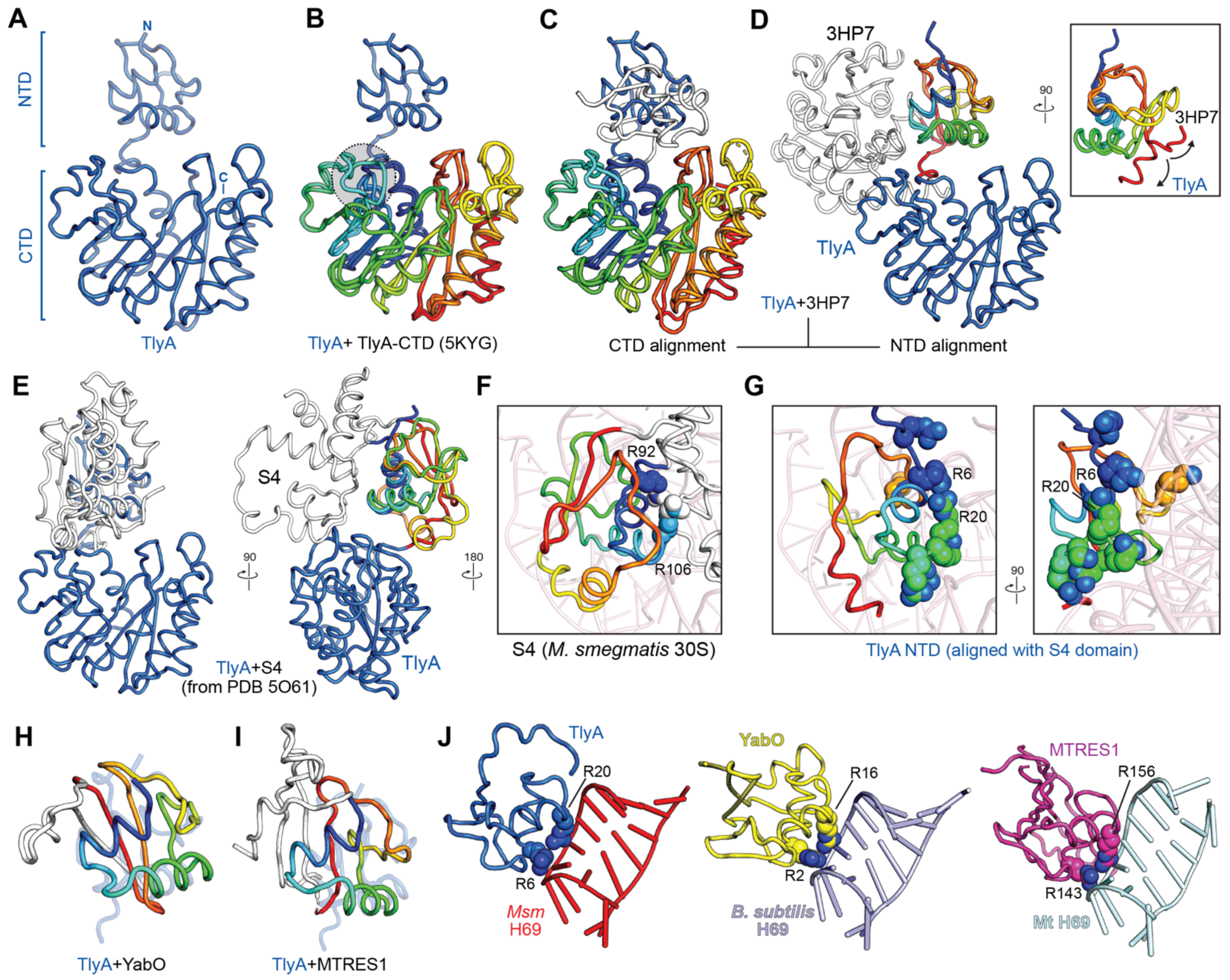


**Fig. S5. Superposition of TlyA with structural homologs**. ***A***, Structure of TlyA with the CTD and *de novo* modeled NTD indicated. ***B***, Superposition of TlyA with the previously determined TlyA CTD crystal structure (PDB code 5KYG). The regions used for alignment are colored as rainbow from N- (blue) to C-terminus (red). Restructuring of the TlyA loop surrounding residue Tyr115 upon 50S subunit binding is highlighted (circle with gray shading). ***C***, Superposition of TlyA with the putative hemolysin from *S. thermophilus* (PDB code 3HP7) aligned based on the protein CTDs (rainbow coloring). ***D***, As *panel C*, but aligned based on the protein NTDs (rainbow coloring). The inset shows a 90˚ rotated view highlighting the different backbone paths taken at the junction of NTD and CTD in TlyA and the putative hemolysin from *S. thermophilus*, which results in the widely differing relative domain orientations. ***E***, Two views of the superposition of TlyA with *Msm* ribosomal protein S4 (from the structure of the complete *Msm* 70S, PDB code 5O61). Alignment was based on the TlyA NTD and corresponding region in S4 (shown in rainbow coloring on the right image). ***F***, View of S4 on the 16S rRNA with residues corresponding to the functionally critical TlyA Arg6 (S4 Arg92) and Arg20 (S4 Arg106) shown as spheres. ***G***, Two views of the TlyA NTD positioned on 16S rRNA by superposition with S4. The residues of TlyA tested for their potential role in 23S rRNA interaction are shown as spheres, including the two functionally critical residues Arg6 and Arg20. ***H***,***I*** Alignment of the TlyA NTD (semi-transparent blue) with ***H***, *Bacillus subtilis* YabO and ***I***, human mitochondrial MTRES1 shown as rainbow coloring for the S4 fold only. ***J***, Comparison of TlyA, YabO and MTRES1 bound to their respective ribosomal RNA H69; TlyA R6 and R20 and corresponding residues in the other proteins are shown as spheres.

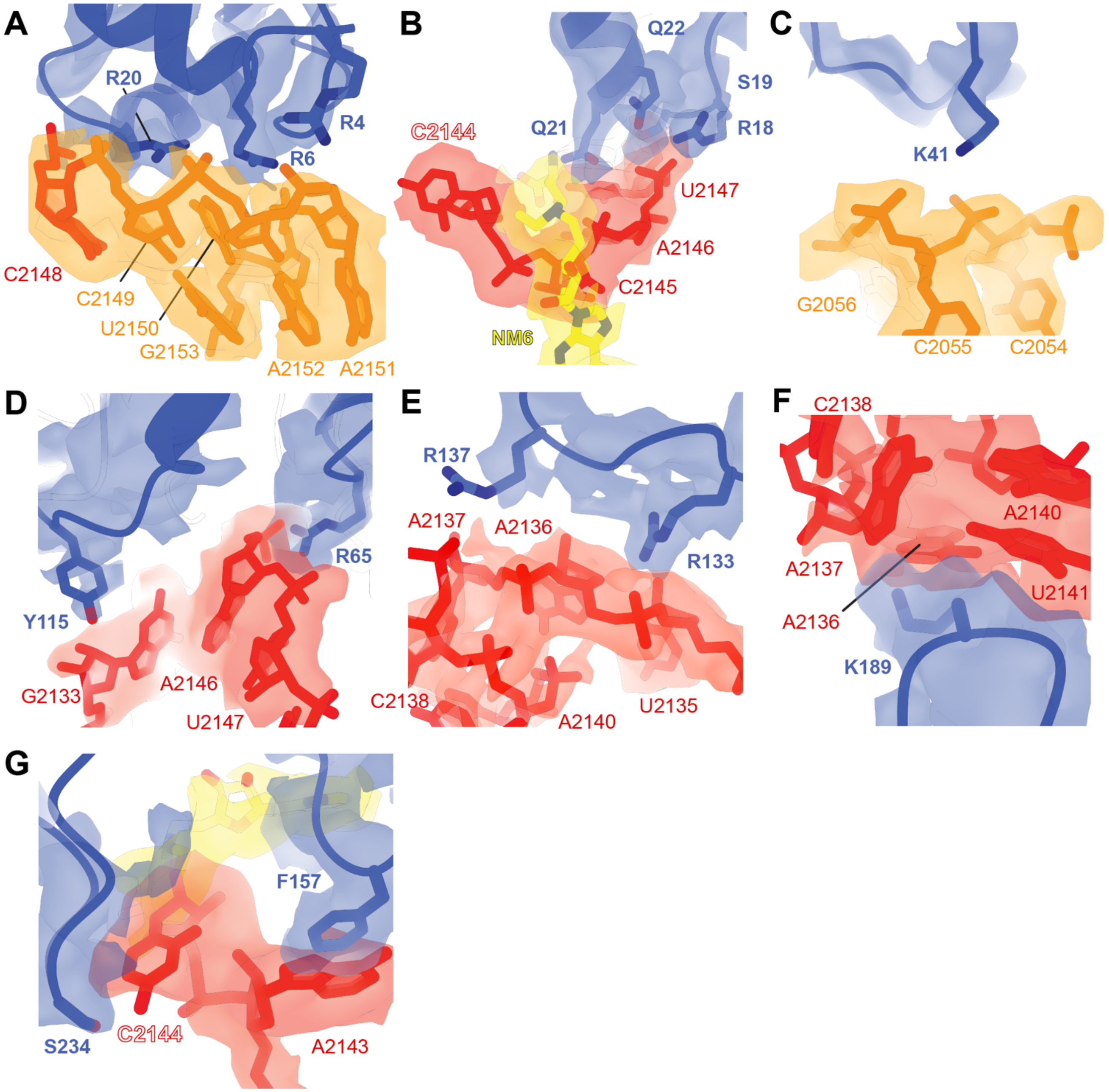


**Fig. S6. Interactions of TlyA NTD and CTD residues with 23S rRNA**. Views corresponding to main figure panels of the TlyA NTD and CTD interactions with H69 (red) and the adjacent rRNA junction (orange) shown with the final complete 50S-TlyA complex map (EMD-24792): ***A,*** TlyA NTD residues Arg4, Arg6 and Arg20 with nucleotides of the rRNA junction proximal to H69 (also see **Fig. 3B**). ***B***, TlyA NTD residues Arg18, Ser19, Gln21 and Gln22 with H69 and NM6 (yellow) (also see **Fig. 3F**). ***C***, TlyA NTD residue Lys41 with nucleotides of the rRNA junction proximal to H69 (also see **Fig. 3G**). ***D***, TlyA CTD residues Arg65 and Try115 with H69 (also see **Fig. 4B**). ***E***, TlyA CTD residues Arg133 and Arg137 with H69 (also see **Fig. 4C**). ***F***, TlyA CTD residue Lys189 with H69 (also see **Fig. 4D**). ***G***, TlyA CTD residues Phe157 and Ser234 with H69 at the site of C2144 base flipping (also see **Fig. 5D**).

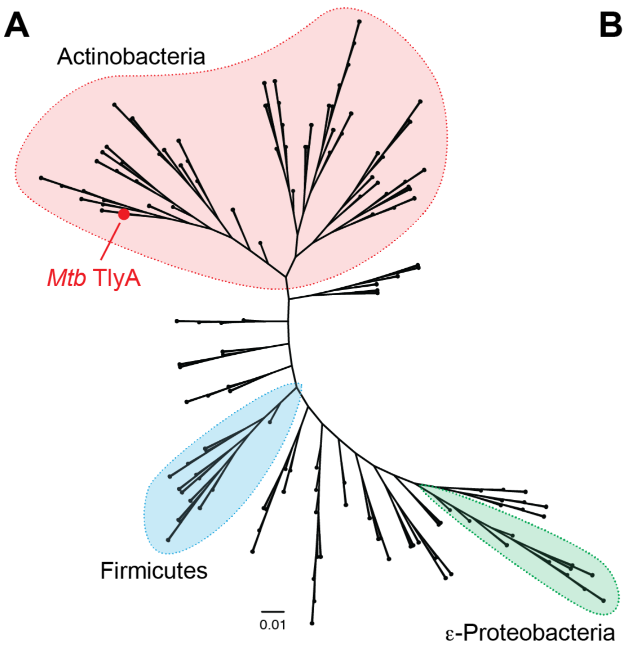


**B**

| TlyA residue | % Conservation (residue identity) | |
|---|---|---|
| | **All homologs** | **Actinobacteria** |
| Arg4 | 74.1 (K/R) | 58.8 (K/R) |
| Arg6 | 97.3 (R) | 100 (R) |
| Arg18 | 15.5 (R)/ 35.5 (D/E) | 34.0 (R)/ 30.9 (D/E) |
| Ser19 | 78.6 (S),15 (T) | 85.6 (S), 9.3 (T) |
| Arg20 | 90.0 (R) | 95.9 (R) |
| Gln21 | 14.3 (S), 20.2 (T) | 16.5 (S), 30.9 (T) |
| Gln22 | 13.0 (Q), 54.7 (K/R) | 13.4 (Q), 50.6 (K/R) |
| Lys41 | 88.3 (K) | 85.4 (K) |
| Arg65 | 97.8 (R) | 100 (R) |
| Tyr115 | 35 (Y), 28.7 (H) | 16.5 (Y), 55.7 (H) |
| Arg133 | 41.3 (R), 14.3 (K) | 30.9 (R), 10.3 (K) |
| Arg137 | 84.8 (R) | 90.7 (R) |
| Phe157 | 97.8 (F) | 99 (F) |
| Lys189 | 51.6 (R), 23.3 (K) | 57.7 (R), 19.6 (K) |
| Ser234 | 11.5 (S), 38.7 (D/E) | 15.6 (S), 47.9 (D/E) |

**Fig. S7. Phylogenetic analysis of the TlyA enzyme family and residue conservation. *A*,** Phylogenetic tree of TlyA homologs inferred using the neighbor-joining method, identifying a major Actinobacterial clade (red) which includes the *M. tuberculosis* TlyA, and two smaller clades with TlyA/ hemolysin homologs from the Firmicutes and Proteobacteria. The scale bar denotes the number of amino acid substitutions per site. ***B***, Residue conservation at sites targeted for mutagenesis in this work. The predominant residue(s) as percentage are shown for TlyA proteins among all homologs or in actinobacteria only.

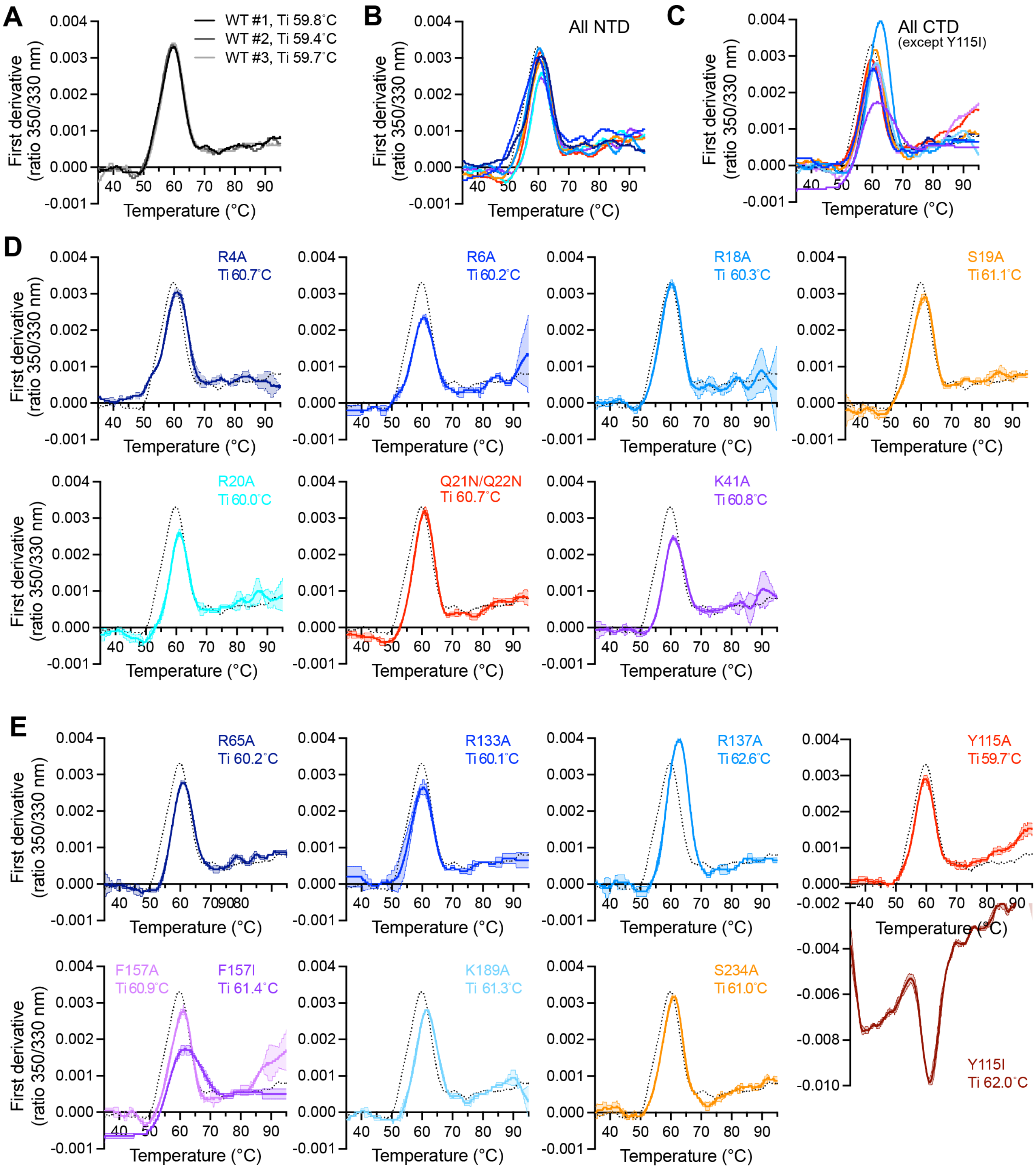


**Fig. S8**. **Quality control of purified wild-type and variant TlyA proteins by thermal denaturation.** ***A***, Replicate measurements of wild-type (WT) TlyA unfolding monitored using intrinsic fluorescence at 330 and 350 nm (Tycho analysis), illustrating the reproducibility of the method between experiments and protein preparations. First derivative plots are shown for fluorescence ratio (350/330 nm) from which inflection temperatures ($T_i$) are derived and shown above the plot. Equivalent analysis for: ***B***, all NTD variants, ***C***, all CTD variants (except Y115I), ***D***, individual NTD variants, and ***E***, individual CTD variants.

Wild-type TlyA is shown for comparison in all panels (black dotted line; average of all measurements in *panel A*). The shaded regions on individual curves show the standard deviation between measurements.

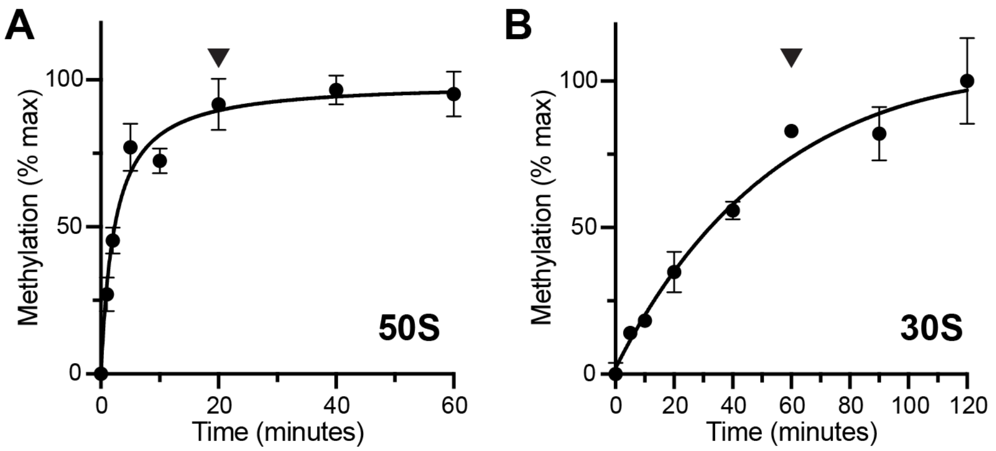


**Fig. S9. Optimization of a [$^{3}$H]-SAM methyltransferase assay for TlyA variants.** Time-course activity assays for wild-type TlyA methylation of ***A***, 50S or ***B***, 30S ribosomal subunits isolated from a *Msm* strain lacking TlyA activity. Methylation is ~90% complete at 20 minutes under the conditions used for 50S subunit and this time point (marked with an arrowhead) was used for single-time point activity analyses of all TlyA variants in the [$^{3}$H]-SAM methyltransferase assay. As 30S subunit is a poorer substrate (see Methods and Methods), a longer time point of 60 minutes (marked with an arrowhead in *panel B*) was used for all analyses of TlyA variants with this substrate.

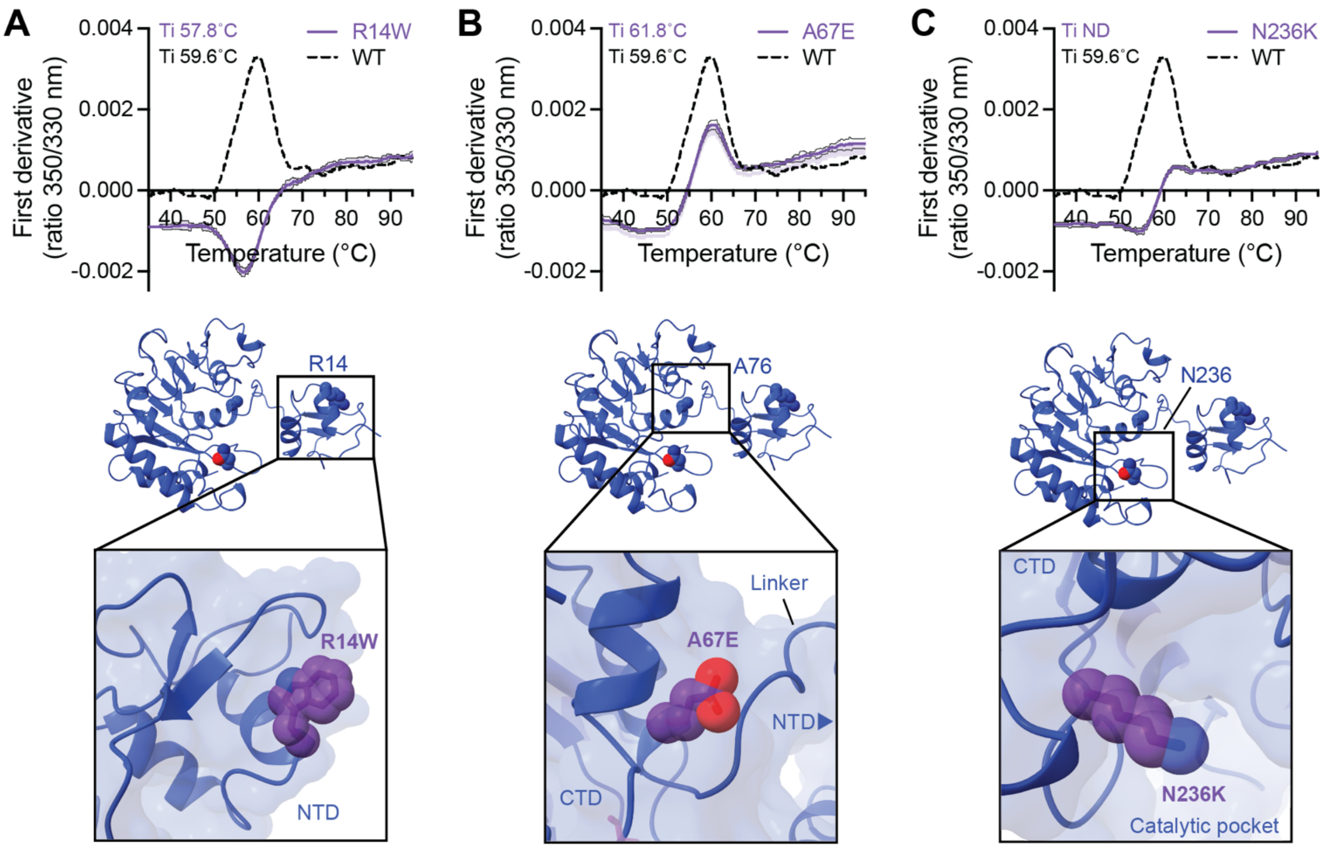


**Fig. S10. Analysis of TlyA clinical variant proteins.** Replicate nDSF measurements of TlyA variant unfolding for: ***A***, R14W, ***B***, A67E and ***C***, N236K, monitored using intrinsic fluorescence at 330 and 350 nm (Tycho analysis). First derivative plots are shown for fluorescence ratio (350/330 nm) from which inflection temperatures ($T_i$) are derived and shown above the plot (ND, not determined). Wild-type TlyA is shown for comparison in all panels (black dashed line; same data as shown in **Fig. S8**). The shaded regions on individual curves show the standard deviation between measurements. Below each plot is a view of the TlyA structure and zoomed views highlighting the residue change in each variant (amino acid substitutions were generated in PyMol).
